# Claudin5 protects the peripheral endothelial barrier in an organ and vessel type-specific manner

**DOI:** 10.1101/2022.03.14.484246

**Authors:** M Richards, E Nwadozi, S Pal, P Martinsson, M Kaakinen, M Gloger, E Sjöberg, K Koltowska, C Betsholtz, L Eklund, S Nordling, L Claesson-Welsh

## Abstract

The pathogenesis of numerous diseases is characterised by disruption of the junctions that form the endothelial cell (EC) barrier, the composition of which may differ greatly between organs. However, the expression level variability and precise contribution of different junctional proteins is poorly understood. Here, we focus on organs with continuous endothelium to identify structural and functional *in vivo* characteristics of the EC barrier. Assembly of multiple single-cell RNAseq datasets into a single integrated database revealed the variability in EC barrier patterning. Across tissues Claudin5 exhibited diminishing expression along the arteriovenous axis, which correlates with EC barrier integrity. Functional analysis identified tissue-specific differences in leakage patterning and response to agonist-induced leakage. We uncover that Claudin5 loss enhances agonist-induced leakage in an organotypic, vessel type-specific and size-selective manner in an inducible, EC-specific, knock-out mouse. Mechanistically, Claudin5 loss induces no change in junction ultrastructure but alters composition, with concomitant loss of zonula occludens-1 (ZO-1) expression and upregulation of VE-Cadherin. These findings uncover the organ-specific organisation of the EC barrier and distinct importance of Claudin5 in different vascular beds and will aid our ability to modify EC barrier stability in a targeted, organ-specific manner.

## Introduction

Blood vessel dysfunction is a hallmark and contributing factor to the progression of numerous pathologies including solid tumours, ischemic diseases and inflammatory conditions (Lee et al., 2007, Burke and Miles, 1958, Hashizume et al., 2000). In such pathologies inflammatory cytokines and growth factors are released, leading to weakening of the endothelial cell (EC) barrier, a regulatable interface between circulating blood and the surrounding tissue environment (Senger et al., 1983, Miles and Miles, 1952, Palade et al., 1979). Loss of EC barrier integrity in-turn leads to enhanced molecular and cellular passage across the endothelium resulting in edema, tissue damage and atrophy and disease progression (Wu et al., 2014, Fleckenstein et al., 2018). EC barrier integrity is mediated primarily by adherens junctions (AJs) and tight junctions (TJs), at which transmembrane proteins form intercellular interactions to bridge adjacent EC membranes (Corada et al., 1999, Claesson-Welsh et al., 2021). These transmembrane proteins also associate with intracellular scaffolding and signalling proteins, as well as the cytoskeleton, which regulate their localisation and stability. The composition and organisation of AJs and TJs thus determine the relative strictness and regulatable nature of the EC barrier.

AJs are ubiquitously distributed in all tissues and vessel subtypes and consist of the largely endothelial-specific transmembrane protein vascular endothelial (VE)-Cadherin, which forms homophilic interactions between cells and associates intracellularly with the actin cytoskeleton via several members of the catenin family (Bazzoni and Dejana, 2004). VE-cadherin-catenin interactions are highly regulated by their phosphorylation status, which modulates AJ stability and integrity of junctions (Smith et al., 2020, Eliceiri et al., 1999, Orsenigo et al., 2012).

TJs are also ubiquitously distributed but are in comparison relatively less well defined, and their composition and localisation appears more heterogeneous in comparison AJs. Numerous transmembrane proteins including members of the claudin, tight junction-associated MARVEL proteins, such as Occludin, junction-associated molecule (JAM) families and endothelial-cell selective adhesion molecule (ESAM) are associated with TJs (Greene et al., 2019, Nasdala et al., 2002, Martin-Padura et al., 1998, Raleigh et al., 2010). With few exceptions, TJ proteins are not specifically expressed by vascular ECs. Similar to AJs, TJ proteins exist in homophilic complexes between adjacent ECs, as well as with intracellular scaffolding proteins including members of the zonula occludens (ZO) family and cingulin (Stevenson et al., 1986, Schossleitner et al., 2016). The relative proportion of these proteins within junctions is believed to influence barrier strictness, however little is known about the actual composition of TJs and how each component contributes to EC barrier integrity, particularly *in vivo*.

Of the TJ proteins Claudin5 is the best studied regarding its contribution to EC barrier integrity. Claudin5 is highly expressed in the brain vasculature and is an important component of the blood brain barrier (BBB) and blood retinal barrier (BRB) where it is responsible for restricting the passage of small molecules (Daneman et al., 2010). Consequently, constitutive *Cldn5* knock-out in mice results in aberrant BBB permeability and death 10 hours after birth (Nitta et al., 2003). Furthermore, downregulation of Claudin5 expression, and resulting enhanced EC permeability, is associated with neurological disorders such as multiple sclerosis, stroke, epilepsy and schizophrenia (Alvarez et al., 2011, Knowland et al., 2014, Yan et al., 2018, Sun et al., 2004). Outside of the CNS, Claudin5 is expressed to a lower degree, as are other TJ proteins such as Occludin (Scalise et al., 2021). Consequently, blood vessels outside of the CNS have a more permeable EC barrier, although EC permeability also differs greatly between these peripheral tissues and is inherently linked to the organ-specific function (Richards et al., 2021, Augustin and Koh, 2017). Furthermore, EC barrier integrity differs between vessel subtypes within these vascular beds (McDonald, 1994, Honkura et al., 2018). Typically the arteriolar aspect of the vasculature is more resistant to EC barrier disruption than the venular side. Interestingly, at least in the mouse ear dermis, EC barrier integrity correlates with Claudin5 expression, which is evident in arterioles but progressively diminishes in capillaries and venules (Honkura et al., 2018). Claudin5 expression and patterning is however poorly understood in tissues outside the CNS, as is the arrangement of other TJ proteins. Furthermore, little is known about the differential impact of TJ components on EC barrier integrity outside of the CNS. Better understanding of EC barrier composition and development of tools to both open and close the barrier at will is an important goal in clinical medicine, as specific manipulation of the EC barrier would allow alleviation of pathogenesis in a range of diseases and improved drug delivery.

In this study analysis of a single-cell RNAseq (scRNAseq) datasets from multiple peripheral vascular beds demonstrates that in general EC junction proteins have a similar distribution in tissues with a continuous endothelium, although with subtle variability in TJ patterning. In all tissues examined here; ear skin, back skin, trachea, skeletal muscle and heart, *Cldn5* exhibits decreasing expression along the arteriovenous axis. In contrast, other TJ components are more broadly expressed in all vessel subtypes. We determined that *Cldn5* expression inversely correlates with histamine-induced vascular leakage, although this patterning exhibits tissue-specific variability. Meanwhile, analysis of Claudin5 function in adult mice using an inducible, endothelial-specific, knock-out mouse also showed organotypic differences, with weakening of the EC barrier occurring to different degrees and in a size-selective manner in distinct tissues. Ultrastructurally, loss of Claudin5 had no effect on EC junction organisation but led to changes in the expression of other junction proteins including VE-Cadherin, ZO-1 and Occludin. Together these data uncover the organotypic heterogeneity that exists in the organisation of EC junctions and the importance of TJ components for general EC barrier integrity.

## Results

### Patterning of the EC barrier at the single-cell level

The organotypic properties of EC junctions were first addressed by exploring the expression profiles of junctional genes in publicly available murine scRNAseq datasets of heart, skeletal muscle and tracheal blood vascular ECs (BECs) (Kalucka et al., 2020, Tabula Muris, 2020) in combination with newly generated scRNAseq data of mouse dermal BECs (Supplemental figure 1). In order to ensure maximum comparability between vessel subsets (i.e. arterial, capillary and venous) in different organs these datasets were integrated using mutual nearest neighbor (MNN) alignment to correct for differences between similar populations, followed by trajectory inference using the tSpace algorithm (Figure 1A) (Haghverdi et al., 2018, Dermadi et al., 2020). An isolated trajectory spanning from arterial to venous BECs of the integrated data was subjected to equidistant binning (Figure 1B) (Dermadi et al., 2020) and the expression of vessel subset-specific markers were subsequently used to guide the annotation of the bins as arterial, arterial/capillary, capillary, capillary/venous, or venous (Figure 1C and D) (He et al., 2018, Vanlandewijck et al., 2018, Kalucka et al., 2020, Brulois et al., 2020). Allocation of the designated subsets was validated by standalone analysis of each individual organ and by comparison to previously published cluster annotations (Supplemental figure 2). This analysis provides a comprehensive integrated database of gene expression across comparable BEC subsets in multiple tissues.

**Figure 1:**
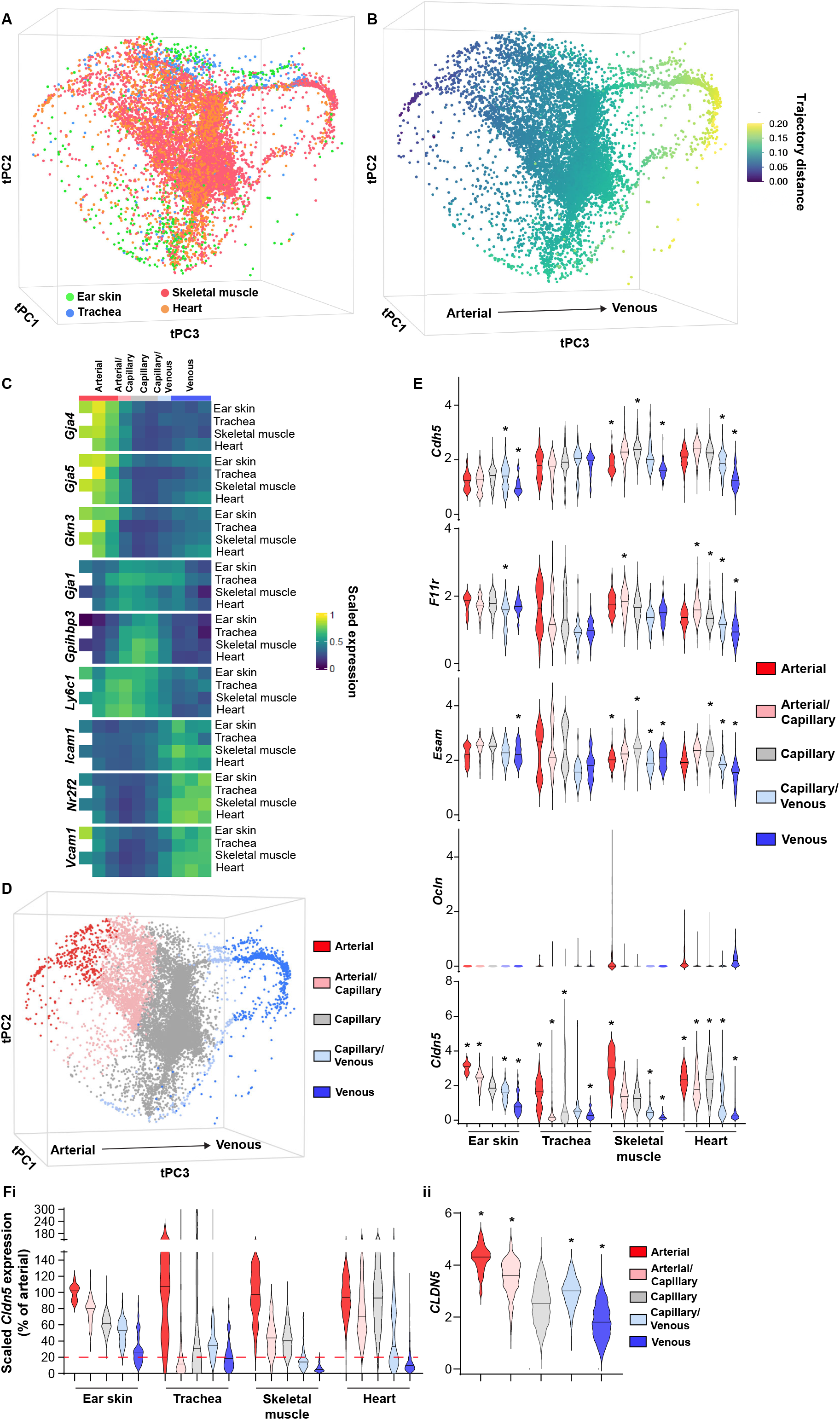
Patterning of the EC barrier at the single cell level. **A**. Principal component analysis of the distances within 400 trajectories calculated with integrated data of murine datasets of ear skin, trachea, skeletal muscle, and heart blood endothelial cells BECs. Colours illustrate the distribution of BECs for each organ. **B**. Principal component analysis of trajectory distances coloured by the distance along an isolated trajectory spanning from arterial to venous BEC. **C**. Mean gene expression for each organ after equidistant binning of the isolated trajectory shown in B. Supervised vessel subset specifications (Top) based on the expression of previously established marker genes. **D**. Principal component analysis of trajectory distances coloured by the vessel subsets defined in C. **E**. Violin plots of gene expression for BEC junctional components. Gene expression was normalized to account for differences in sample library size and has been imputed to account for dropouts in the data as described in Methods. **F**. **i.** *Cldn5* expression in murine BEC datasets scaled per organ according to the mean expression in the arterial vessels of each organ. Red dashed line represents a 5-fold reduction in expression compared to arterial ECs. **ii.** *CLDN5* expression in human dermal BECs. n = 534 ear skin, 559 trachea, 3498 skeletal muscle, 6423 heart and 8518 human BEC. **\*** denotes statistical significance following differential gene expression analysis (Supplemental data 1 and 2).

Subsequently, the expression of genes associated with EC junctions was investigated in each organ and vessel subset (Figure 1E, Supplemental figure 3). A differential gene expression analysis between each subset and all other cells in an organ was utilized to guide conclusions (Supplemental data 1). The adherens junction gene *Cdh5* (VE-Cadherin) was ubiquitously expressed within each organ but was generally slightly lower in venous vessels compared to other subsets within these tissues (Figure 1E). Other Cadherins such as N-Cadherin (*Cdh2*) and T-Cadherin (*Cdh13*) were evenly expressed across vessel subsets, although at low and relatively high levels respectively (Supplemental figure 3). Tight junction genes displayed differing expression patterns. The JAM family of junctional adhesion molecules (*F11r, Jam2, Jam3*) exhibited reasonably homogenous expression, but were occasionally more abundantly expressed in capillaries than in arteries and veins, in an organotypic manner (Figure 1E and Supplemental figure 3). *Esam* similarly was homogenously expressed with reduced expression in the venous subset of some tissues (Figure 1E). Expression of the tight junction-associated Marvel proteins (TAMPs) *Ocln, Marveld2* and *Marveld3* meanwhile were relatively low, being limited to only a few BECs in an organotypic and subset-specific manner (Figure 1E and Supplemental figure 3). Most members of the Claudin family were expressed at a very low level with the exception of *Cldn5* and *Cldn15*. Both these genes exhibited diminishing expression along the arteriovenous axis, with *Cldn5* in particular possessing higher expression in arteries compared to veins, displaying fold changes between 3 (ear skin) and 15 (skeletal muscle) (Figure 1E and Fi). Accordingly, analysis of genes differentially expressed between subsets showed consistently significant changes in *Cldn5* expression (Supplemental data 1).

We further utilized a recently published human skin BEC scRNAseq dataset to investigate whether EC junctional patterning is faithfully conserved between human and mouse (Li et al., 2021). Similar analysis of human skin BECs, as employed for mouse BECs, showed enrichment of adherens junction genes *CDH5* and *CDH13* in the arterial and capillary subsets, with lower levels in venous proximal capillaries and veins (Supplemental figure 4 and 5). In keeping with the overall trend in the mouse vasculature, human tight junction component expression was also relatively higher in arteries and/or capillaries; *F11R, ESAM, CLDN5, CLDN10*, and *CLDN15* expression was significantly higher in arteries, while *ESAM, CLDN5*, and *CLDN15* were downregulated in veins compared to all other subsets (Figure 1Fii, Supplemental figure 5 and Supplemental data 2).

### Organotypic regulation of barrier integrity

In the ear dermis, expression of Claudin5 inversely correlates with vessel susceptibility to Vascular Endothelial Growth Factor A (VEGF-A)-induced leakage (Honkura et al., 2018). We thus decided to study whether EC barrier integrity was similarly patterned in other tissues and in response to other agonist classes such as inflammatory cytokines. For this purpose, we employed histamine in an intravital imaging setup coupled with atraumatic intradermal agonist injection, in which acute leakage may be accurately assessed. Claudin5 expression was followed through *Cldn5* promoter-driven expression of GFP (*Cldn5*(BAC)-GFP) (Honkura et al., 2018). Here, intradermal administration of histamine elicited a similar leakage patterning as previously shown for VEGF-A, with leakage occurring only in vessels lacking apparent *Cldn5*(BAC)-GFP expression (Figure 2Ai and ii and Movie 1). Interestingly, a population of *Cldn5*(BAC)-GFP-negative vessels that failed to respond to histamine stimulation exists, highlighting the potential importance of factors other than Claudin5 that influence vessel susceptibility to stimulation (Figure 2Aiii) (Richards et al., 2021).

**Figure 2:**
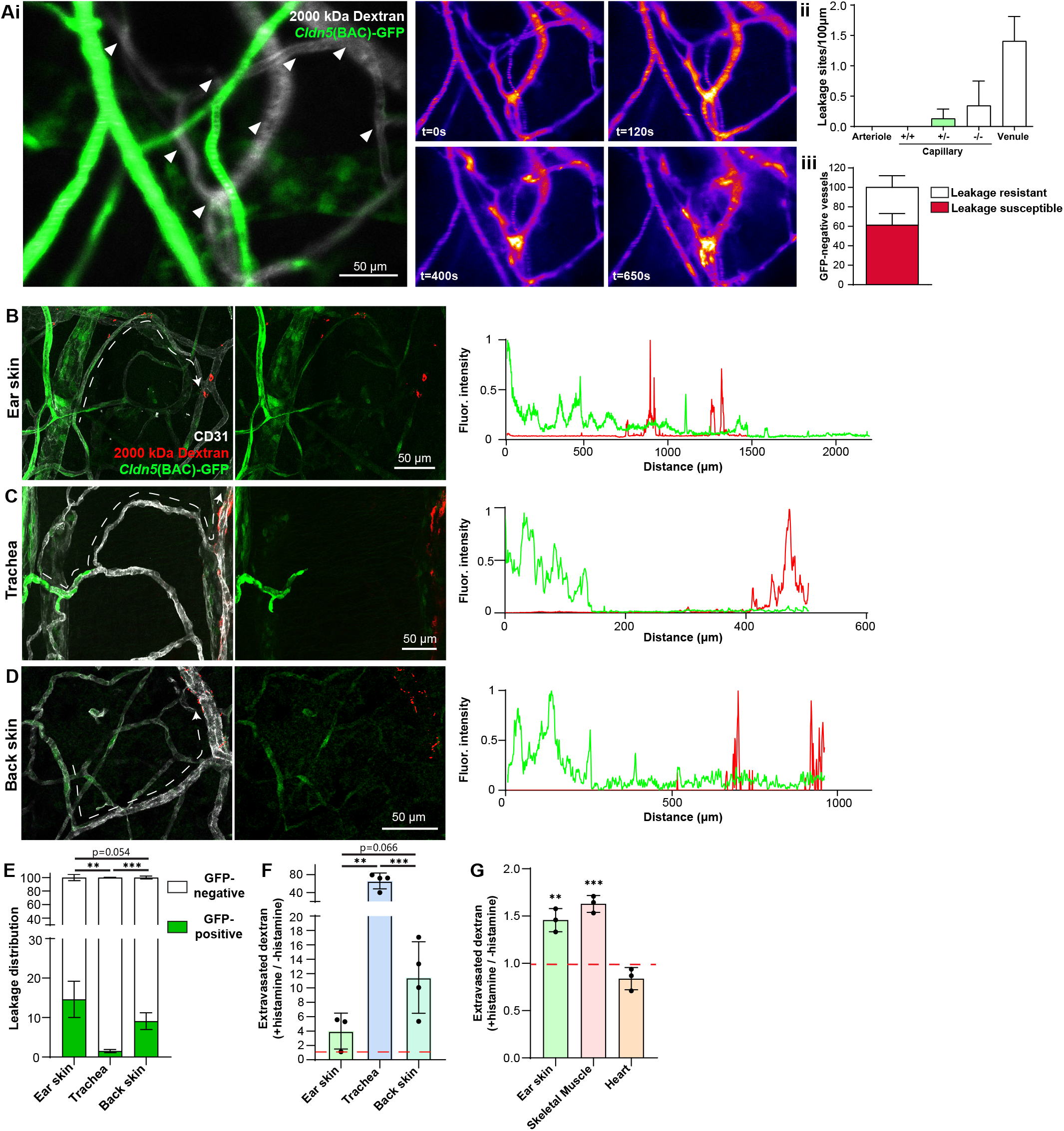
Organotypic integrity of the EC barrier. **A. i.** Leakage patterning in *Cldn5*(BAC)-GFP mouse ear skin in response to intradermal histamine. Left, overlay of *Cldn5*(BAC)-GFP-positive and -negative vessels (visualised through circulating TRITC dextran). Arrowheads show sites of leakage. Right, Stills of vascular leakage of the vasculature shown on the left following intradermal histamine stimulation. **ii.** Leakage sites per vessel length in different vessel categories. **iii.** Proportion of *Cldn5*(BAC)-GFP-negative vessels susceptible or resistant to leakage. n = 4, 2 or more acquisitions/mouse. **B.** Leakage patterning in the ear skin in response to the systemic administration of histamine. Left, representative image. Dashed line shows progression of a blood vessel from arteriolar to venular. Right, representative fluorescent intensity line profile of *Cldn5*(BAC)-GFP and TRITC 2000 kDa dextran along the dashed line (Left). **C.** Leakage patterning in the trachea in response to the systemic administration of histamine. Left, representative image. Dashed line shows progression of a blood vessel from arteriolar to venular. Right, representative fluorescent intensity line profile of *Cldn5*(BAC)-GFP and TRITC 2000 kDa dextran along the dashed line (Left). **D.** Leakage patterning in the back skin in response to the systemic administration of histamine. Left, representative image. Dashed line shows progression of a blood vessel from arteriolar to venular. Right, representative fluorescent intensity line profile of *Cldn5*(BAC)-GFP and TRITC 2000 kDa dextran along the dashed line (Left). **E.** Proportion of 2000 kDa FITC leakage area that occurs in vessels that are *Cldn5*(BAC)-GFP-positive (contain some positive cells) and *Cldn5*(BAC)-GFP-negative (contain no positive cells) in ear skin, back skin and trachea. n ≥ 3 mice, 3 or more fields of view/mouse. **F.** Fold change in 2000 kDa TRITC dextran extravasation from leakage permissive vessels in ear skin, back skin and trachea with and without systemic histamine stimulation. Dashed line represents unstimulated tissue. n = 3 mice, 3 or more fields of view/mouse. **G.** Fold change in tissue 2000 kDa FITC dextran following systemic histamine stimulation and formamide extraction of ear skin, skeletal muscle and heart. Dashed line represents unstimulated tissue. n = 3 mice. Error bars; mean ± SEM. Statistical significance: two-tailed paired Student’s t test.

Next, a model of acute systemic leakage was used whereby histamine, when injected into the circulation via the tail-vein, results in widespread disruption of BEC junctions and leakage fluorescent tracers (Richards et al., 2021). When given systemically, along with a 2000 kDa lysine-fixable dextran, histamine caused leakage in the ear dermis from venules and also from capillaries exhibiting a salt and pepper expression of *Cldn5*(BAC)-GFP, similar to local administration (Figure 2B). Skeletal muscle also showed leakage susceptibility starting in the capillary bed where *Cldn5*(BAC)-GFP expression was heterogeneous (Supplemental figure 6A). The heart vasculature meanwhile showed no extravascular dextran accumulation with or without histamine stimulation (Supplemental figure 6A). Unlike the ear dermis and skeletal muscle, the tracheal vasculature displayed a clear separation between *Cldn5*(BAC)-GFP-positive arterioles and leakage susceptible vessels, with the capillaries that cross the cartilage rings being *Cldn5*(BAC)-GFP-negative and resistant to stimulation (Figure 2C). Interestingly, examination of the mouse back skin revealed an intermediary phenotype, with vessels possessing a salt and pepper expression of *Cldn5*(BAC)-GFP being largely resistant to leakage, whilst immediately subsequent venules exhibited a strong leakage phenotype (Figure 2D). Representative line profiles demonstrate this variable patterning, with extravasated dextran interspersed with a *Cldn5*(BAC)-GFP-positive signal in the ear dermis (Figure 2B) but not in the back skin or trachea (Figure 2C and D). These differing leakage patterns are also apparent from the increased proportion of extravasated dextran from *Cldn5*(BAC)-GFP-positive vessels in the ear skin compared to back skin and trachea (Figure 2E). Additionally, these tissues differ in their magnitude of leakage to histamine stimulation (Figure 2F). In response to the same stimulus leakage permissive vessels in the trachea and the back skin showed much more extensive leakage as compared to the ear skin. Skeletal muscle showed a similar leakage response as the ear skin whilst, as expected, no histamine-induced leakage was observed in the heart (Figure 2G).

This data shows that outside of the CNS, the zonation of Claudin5 protein expression along the arteriovenous axis varies between organs, although all organs show diminished expression towards the venous end. Moreover, the relationship between Claudin5 expression and susceptibility to vascular leakage differs between vascular beds, in particular in the salt-and-pepper capillary bed, as does the sensitivity of ECs to stimulation and their resulting junctional disruption.

### Claudin5 exhibits organotypic protection of the EC barrier

Claudin5 is a well-known determinant of blood vessel barrier integrity in the CNS. Constitutive Claudin5 deficiency results in the passage of small molecules from the blood into the cerebrospinal fluid and death 10 hours after birth (Nitta et al., 2003). Nothing is known about the role that Claudin5 plays in barrier stability outside of the CNS. To investigate this, we used an inducible, EC-specific *Cldn5* loss-of-function mouse model (*Cldn5* fl/fl; *Cdh5* CreER^T2^ (*Cldn5* iECKO)) (Supplemental figure 6B). Systemic administration of tamoxifen to these mice led to no obvious physiological or behavioural defects the first five days following treatment completion. Analysis of lung lysates showed a near total loss of *Cldn5* RNA (Figure 3Ai) and a 60% reduction in Claudin5 protein levels in iECKO mice compared to controls (Figure 3Aii). Blood vessels in the ear dermis meanwhile showed an approximately 75% reduction in Claudin5 levels, as judged from immunofluorescent staining, with some remaining protein expression evident in arterioles (Figure 3B). In an attempt to enhance the loss of Claudin5 without causing lethality, mice were treated topically on the ear skin with 4-hydroxytamoxifen, followed by a 14-day interval before tissue collection. Similar to systemically treated mice, Claudin5 loss was limited to approximately 75% (Figure 3Bii). The use of a Rosa26 STOP fl/fl YFP reporter revealed that in arterioles Claudin5 protein levels remain even though recombination has occurred (Figure 3C), suggesting that there is a pool of junctional Claudin5 that is extremely stable and subject to slower turnover.

**Figure 3:**
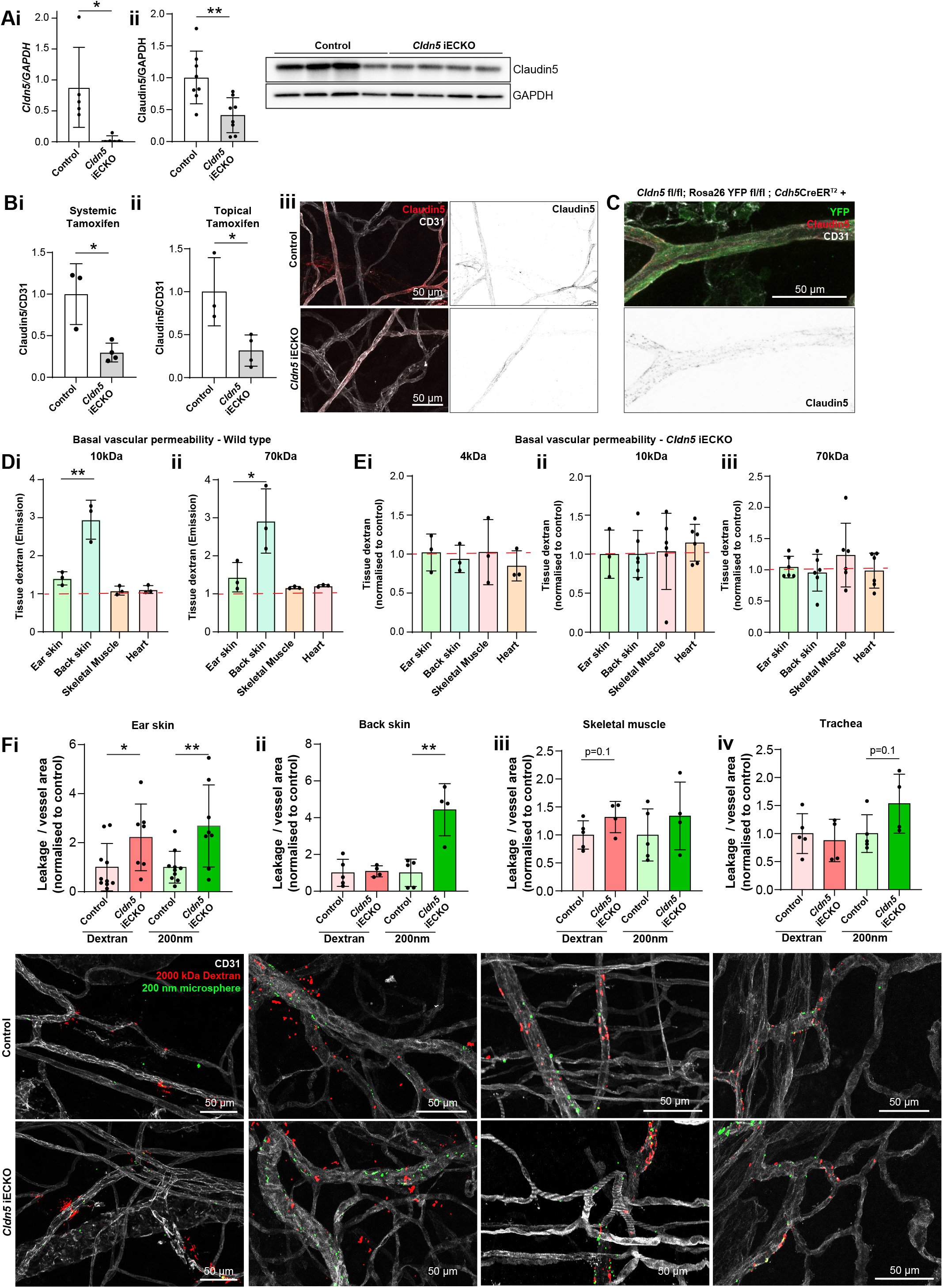
Claudin5 exhibits organotypic protection of the EC barrier. **A. i.** *Cldn5* gene expression by qPCR on lung lysates of control and *Cldn5* iECKO mice. **ii.** Claudin5 protein expression in lung lysates of control and *Cldn5* iECKO mice. Left, quantification. Right, representative blot. n ≥ 5 mice. **B.** Claudin5 protein expression in the ear skin of control and *Cldn5* iECKO mice following systemic (**i**) and topical tamoxifen (**ii**). **iii**. Representative images of Claudin5 immunostaining in control and *Cldn5* iECKO mice. n ≥ 3 mice, 3 or more fields of view/mouse. **C.** Representative image of Claudin5 expression in arterioles of *Cldn5 fl/fl*; Rosa26 YFP fl/fl; *Cdh5*CreER^T2^ mice following tamoxifen treatment. **D.** Blood vessel basal permeability to 10 kDa (**i**) and 70 kDa (**ii**) dextran in ear skin, back skin, skeletal muscle and heart of wildtype C57Bl/6 mice. Dashed lines represent control uninjected mice. n = 3 mice. **E.** Blood vessel basal permeability to 4 kDa (**i**), 10 kDa (**ii**) and 70 kDa (**iii**) dextran in ear skin, back skin, skeletal muscle and heart of control and *Cldn5* iECKO mice. Dashed lines represent control Cre-negative mice. n ≥ 3 mice. **F.** Leakage of 2000 kDa dextran and 200 nm microspheres in response to systemic histamine stimulation in ear skin (**i**), back skin (**ii**), skeletal muscle (**iii**) and trachea (**iv**). Top, quantification of tracer leakage area / vessel area normalised to control (Cre-negative) mice. Bottom, representative images. n ≥ 4 mice, 3 or more fields of view/mouse. Error bars; mean ± SEM. Statistical significance: two-tailed paired Student’s t test.

*Cldn5* iECKO mice were next investigated to assess whether Claudin5 maintains the permeability of blood vessels in tissues outside of the CNS. To study this, mice were systemically injected with differently sized fluorescent dextrans, which were allowed to circulate before mice were perfused and organs collected. Extravasated dextran was subsequently extracted into formamide and measured according to their spectra. Initially, to better understand tissue-specific differences in the EC barrier, organs were assessed for their basal permeabilities (Figure 3D). Interestingly, the back skin showed relatively high basal permeability when compared with ear skin, which showed minor dextran extravasation, while there was no basal leakage from skeletal muscle and heart. Loss of Claudin5 meanwhile did not affect basal permeability when comparing *Cldn5* iECKO with Cre-negative control mice, for all sizes of dextran investigated (Figure 3E).

We next sought to determine whether the loss of Claudin5 leads to changes in histamine-induced macromolecular leakage. In *Cldn5* iECKO mice, the heart vasculature still showed resistance to histamine-mediated leakage (Supplemental figure 6C). In the ear dermis, systemic administration of histamine resulted in an increase in the extravasation of both 2000kDa dextran and 200nm microspheres in *Cldn5* iECKO mice (Figure 3Fi). In contrast, back skin, trachea and skeletal muscle showed no significant increase in 2000kDa dextran extravasation following the loss of Claudin5, although a small increase was seen in skeletal muscle (Figure 3Fii-iv). Interestingly, extravasation of larger 200nm microspheres increased significantly in back skin, whilst a slight increase was seen in trachea, following the loss of Claudin5 expression (Figure 3Fii and iv). These data suggest that, as opposed to the CNS where Claudin5 assembles seamless tight junctions, the relative sparsity of Claudin5 in venous ECs outside of the CNS (see Figure 1F) creates junctional strands that restrict excessive disruption of junctions in response to leakage agonists, the limits of which differ between vascular beds.

Claudin5 thus has a limited role in maintaining baseline EC barrier integrity in blood vessels outside of the CNS but is involved in the protection of vessels against agonist-induced macromolecular leakage in an organotypic and size-selective manner.

### Loss of Claudin5 differentially affects vessel subtypes

The increase in histamine-induced permeability in the ear dermis following the loss of Claudin5 prompted us to address in which vessel type the enhanced barrier breakdown is taking place. In the ear skin, expression of Claudin5 in arterioles was clearly reduced and patchier in the *Cldn5* iECKO mice (Figure 4A). Even so, these vessels remained resistant to histamine-induced leakage as revealed by both systemic administration of histamine and intravital visualisation of vascular leakage following intradermal histamine administration (Figure 4A and B and Movie 2). In keeping with the data shown in Figure 3Fi there was still a significant increase in overall vessel leakage, with more leakage sites being induced per vessel length after histamine stimulation in the *Cldn5* iECKO ear dermis (Figure 4Biii and Movie 2). The extent of barrier disruption at each site of leakage was also found to be increased, with individual leakage sites exhibiting enhanced extravasation of dextran following Claudin5 loss (Figure 4Biv). The rate of endothelial response to stimulation was unchanged, with leakage occurring approximately 3 minutes after stimulation in *Cldn5* iECKO mice and their controls (Figure 4Bv).

**Figure 4:**
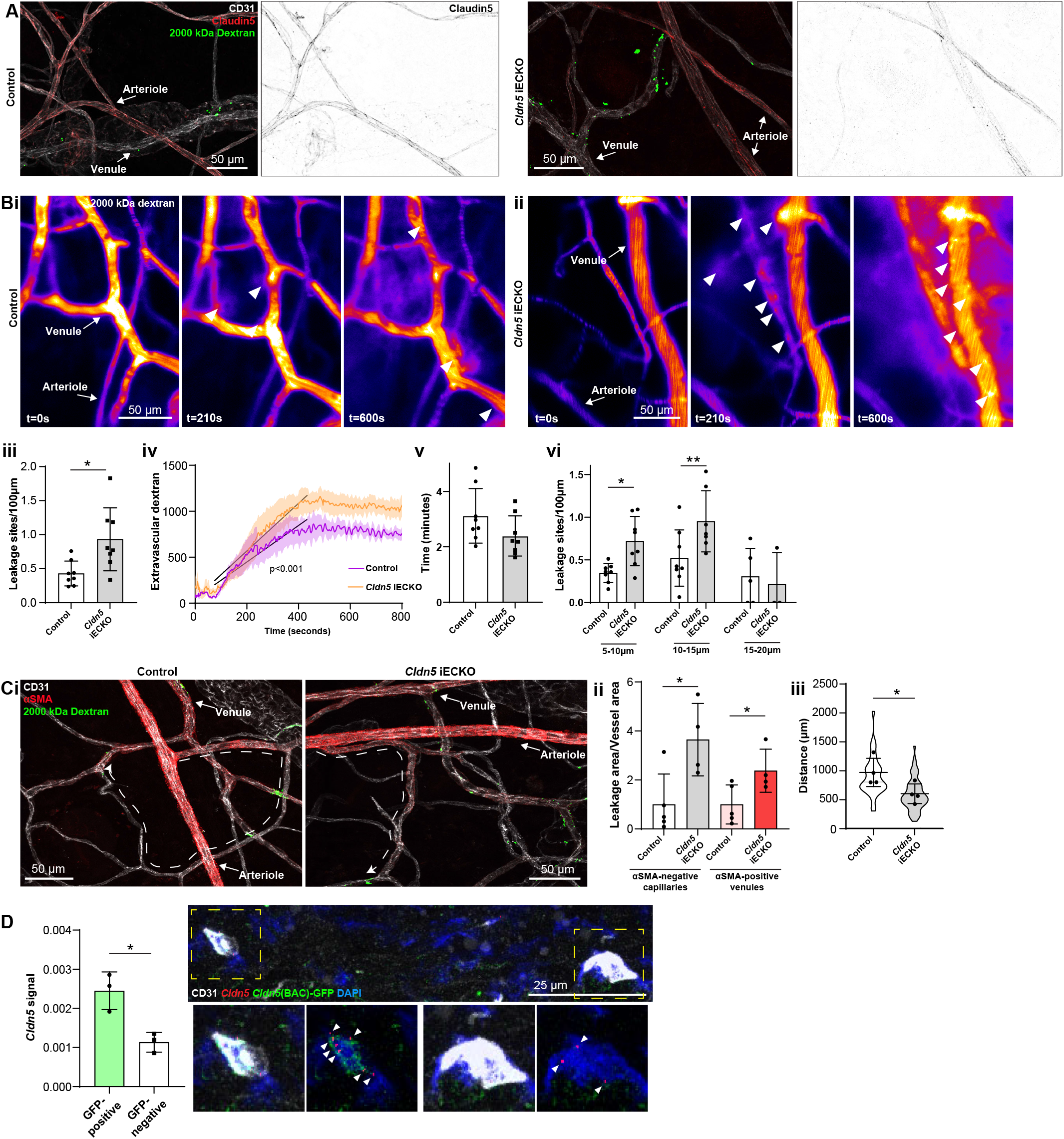
Loss of Claudin5 differentially affects vessel subtypes. **A.** Representative images of histamine-induced 2000 kDa dextran leakage in the ear skin of control (left) and *Cldn5* iECKO (right) mice. **B.** Representative time-lapse images of 2000 kDa dextran leakage in response to intradermal histamine stimulation in the ear skin of control (**i**) and *Cldn5* iECKO (**ii**) mice. Arrowheads show sites of leakage. **iii.** Leakage sites per vessel length in response to intradermal histamine stimulation in the ear skin of control and *Cldn5* iECKO mice. **iv.** Quantification of extravascular 2000 kDa dextran over time in the ear skin of control and *Cldn5* iECKO mice following intradermal histamine stimulation. Black lines represent lines of best fit for the slope between leakage initiation and leakage termination. **v.** Lag period between intradermal histamine injection and initiation of leakage in the ear skin of control and *Cldn5* iECKO mice. **vi.** Leakage sites per length of post-arteriolar vessels of different diameter in response to intradermal histamine stimulation in the ear skin of control and *Cldn5* iECKO mice. n ≥ 7 mice, two or more acquisitions/mouse. **C. i.** Representative images of 2000 kDa dextran leakage in response to systemic histamine stimulation in the ear skin of control and *Cldn5* iECKO mice counter-stained for αSMA. **ii.** Leakage area / vessel area of 2000 kDa dextran in response to systemic histamine stimulation in αSMA-negative capillaries and αSMA-positive venules in the ear skin of control and *Cldn5* iECKO mice. **iii.** Distance between arteriolar-capillary branch points and the first site of 2000 kDa dextran leakage in response to systemic histamine stimulation in the ear skin of control and *Cldn5* iECKO mice. n ≥ 4 mice, 3 or more fields of view/mouse. **D.** *Cldn5* mRNA expression in *Cldn5*(BAC)-GFP-positive and -negative vessels of the ear skin. Left, quantification of *Cldn5* signal (*Cldn5* mRNA particles / vessel area). Right, representative image. Dashed boxes are magnified below, arrowheads mark *Cldn5* mRNA particles. n = 3 mice, 4 or more fields of view/mouse. Error bars; mean ± SEM. Statistical significance: two-tailed paired Student’s t test and linear regression and ANCOVA.

We next wished to explore in which vessel type leakage was enhanced following Claudin5 loss. Segregation of vessels into their subtypes in the dermis is however complicated by their stochastic organisation and clear separation between capillaries, which are often assumed to be leakage resistant, and post-capillary venules (leakage permissive) is challenging. Thus, determining precisely which vessels are affected by loss of Claudin5 is not possible. Post-arteriolar vessels were thus instead separated based on luminal dextran diameter. This analysis showed that small vessels (5-10 μm) and subsequent mid-sized vessels (10-15 μm) leaked to a greater degree following the loss of Claudin5. In contrast, larger venules (15-20 μm) showed no change in leakage (Figure 4Bvi). To explore further where these vessels lie within the vasculature, analysis of systemically-induced histamine leakage was carried out alongside visualisation of α-smooth muscle actin (αSMA), which is absent on capillaries but present on the arteriolar and venular aspect of the microvasculature. In the ear skin of *Cldn5*(BAC)-GFP mice, arteriolar αSMA coverage can be seen to end before *Cldn5*(BAC)-GFP expression, whilst its venular expression begins after *Cldn5*(BAC)-GFP expression has been lost (Supplemental figure 6D). *Cldn5*(BAC)-GFP expression is thus lost in the αSMA-negative capillary bed. Based on this segregation, we observed a 2.5-fold increase in leakage from αSMA-positive venules and a 4-fold increase in leakage in αSMA-negative capillaries in *Cldn5* iECKO mice (Figure 4Ci and ii). Furthermore, in *Cldn5* iECKO mice leakage sites were seen to appear closer to arterioles than in their controls (Figure 4Ciii).

This data demonstrates that loss of Claudin5 decreases the junctional integrity of capillaries and venules, at least in the ear skin, and moves the limit between leakage resistant and susceptible ECs towards the arteriolar aspect. We conclude that loss of Claudin5, to a large degree, enhances histamine-induced leakage in vessels which are Claudin5 negative, according to immunohistochemistry as well as to the GFP reporter mouse used here. Use of more sensitive RNA *in situ* hybridisation however showed that in *Cldn5*(BAC)-GFP-negative vessels *Cldn5* expression still occurs, albeit to a lower level than *Cldn5*(BAC)-GFP-positive vessels, in keeping with the expression patterning observed from scRNAseq data (Figure 1F and Figure 4D).

From the highly sensitive RNAscope detection of *Cldn5* expression in capillaries and postcapillary venules, we conclude that Claudin5 is responsible for limiting the disruption of EC junctions in particular in capillaries but also in immediately following venules. Arterioles remain resistant to agonist-induced leakage, as do larger venules, presumably due to residual Claudin5 expression and the relatively high expression of other TJ components.

### Claudin5 modulates junction protein expression

The consequence of Claudin5 loss was further studied by transmission electron microscopy (TEM), with focus on non-arteriolar vessels in the ear dermis. In TEM the intercellular cleft is lined by parallel plasma membranes of contacting ECs and junctional complexes appear as electron dense structures following uranyl acetate staining. Analysis of the electron dense area, width and density showed that there was no obvious change in this junction structure after loss of Claudin5 (Figure 5Ai-iii). In keeping with the enhanced leakage that we observed following the loss of Claudin5, greater disruption of the endothelial barrier was seen following histamine stimulation, allowing greater penetrance of horse radish peroxidase (HRP) into the intercellular cleft (Figure 5Aiv-v).

**Figure 5:**
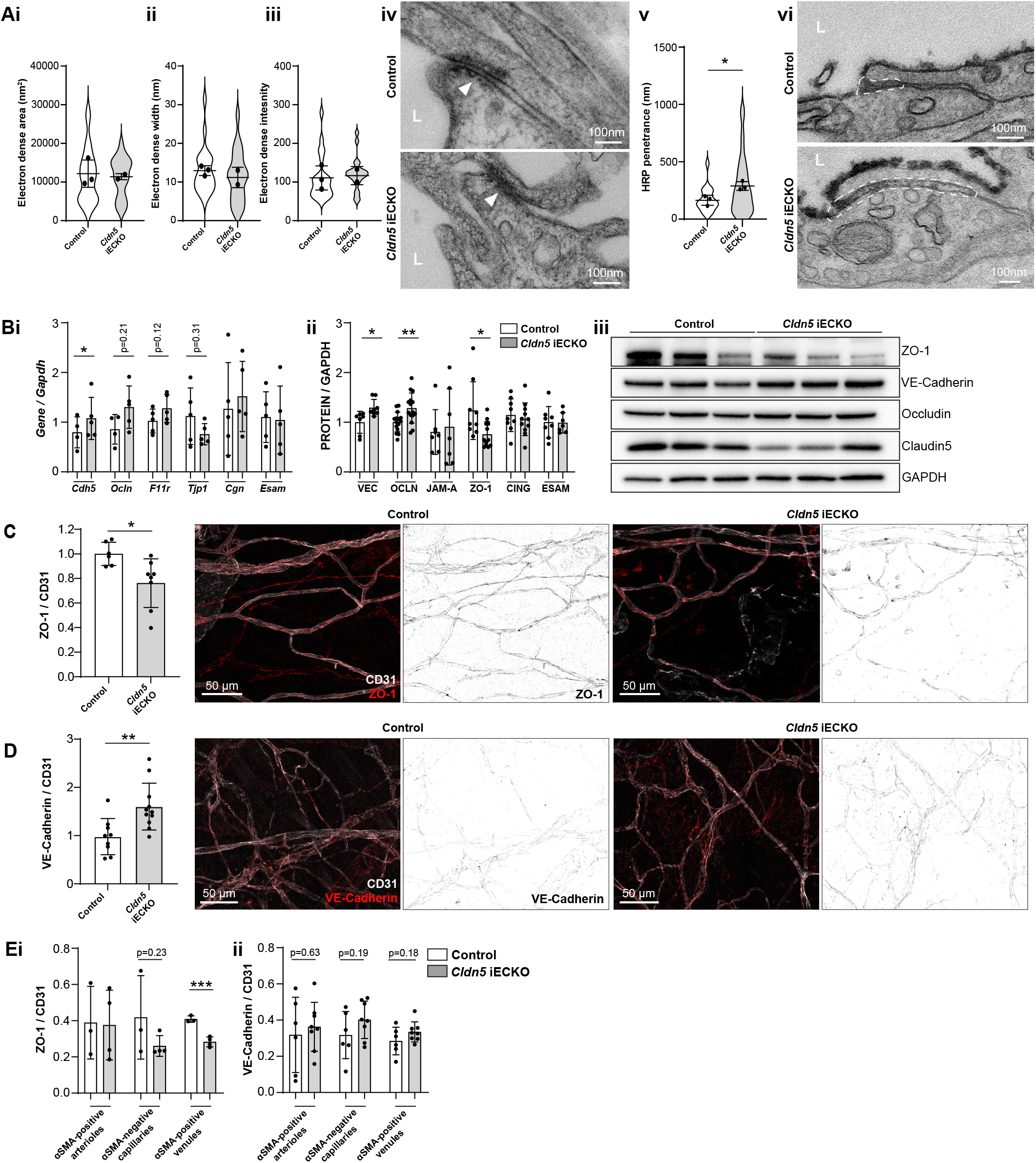
Claudin5 regulates junction protein expression. **A.** Area (**i**) width (**ii**) and intensity (i**ii**) of electron dense regions in the ear skin of control and *Cldn5* iECKO mice after visualisation by TEM. **iv.** Representative TEM images of junctions in the ear skin of control and *Cldn5* iECKO mice. Junctions can be seen within electron dense regions (arrowheads). **iv.** Distance of HRP penetrance into EC junctions in the ear skin of control and *Cldn5* iECKO mice following systemic histamine stimulation. **v.** Representative TEM images of HRP penetrance (visualised by electron dense 3,3′-Diaminobenzidine (DAB) reaction precipitate) into EC junctions in the ear skin of control and *Cldn5* iECKO mice following systemic histamine stimulation. Dashed regions show areas of disrupted junction into which HRP has penetrated. Note that the typical electron dense area is lacking due to absence of uranyl acetate staining. L, lumen. n ≥ 2 mice, 6 or more fields of view/mouse. **B. i.** Gene expression of AJ- and TJ-associated genes in lung lysates of control and *Cldn5* iECKO mice. **ii.** Expression of AJ- and TJ-associated proteins in lung lysates of control and *Cldn5* iECKO mice. **iii.** Representative western blots of AJ- and TJ-associated proteins in lung lysates of control and *Cldn5* iECKO mice. n ≥ 4 mice. **C.** Expression of ZO-1 in ear skin blood vessels of control and *Cldn5* iECKO mice. Left, quantification of ZO-1. Right, representative images of ZO-1 in the ear skin of control and *Cldn5* iECKO mice. n ≥ 6 mice, 3 or more fields of view/mouse. **D.** Expression of VE-Cadherin in ear skin blood vessels of control and *Cldn5* iECKO mice. Left, quantification of VE-Cadherin. Right, representative images of VE-Cadherin in the ear skin of control and *Cldn5* iECKO mice. n ≥ 9 mice, 3 or more fields of view/mouse. **E. i.** Quantification of ZO-1 in different vessel subtypes in the ear skin of control and *Cldn5* iECKO mice. **ii.** Quantification of VE-Cadherin in different vessel subtypes in the ear skin of control and *Cldn5* iECKO mice. n ≥ 3 mice, 3 or more fields of view/mouse. Error bars; mean ± SEM. Statistical significance: two-tailed paired Student’s t test.

The lack of apparent change in junction structure, but change in junction integrity, is surprising but may be explained by compensatory changes in junction composition. It is known that Claudin5 expression, through transcriptional cross-talk, can be controlled through VE-Cadherin and JAM-A (Taddei et al., 2008, Kakogiannos et al., 2020). Whether Claudin5 expression reciprocally controls the expression of other junctional proteins is unknown. We therefore investigated the expression of cell-cell adhesion proteins in *Cldn5* iECKO mice. Initially, lung RNA was screened for the expression of the AJ gene *Cdh5* (VE-Cadherin) and common TJ genes *Tjp1* (ZO-1), *Ocln* (Occludin), *F11r* (JAM-A), *Cgn* (Cingulin) and *Esam* (ESAM). Following *Cldn5* KO small increases in *Cdh5, Ocln* and *F11r* expression, but a decrease in *Tjp1* expression was observed (Figure 5Bi). Subsequent analysis of lung protein samples confirmed the enhanced protein expression of VE-Cadherin and Occludin and decrease in ZO-1 (Figure 5Aii and iii and Supplemental figure 7A). Correlation analysis of these samples also supported this finding, with Claudin5 expression levels inversely correlating with VE-Cadherin and Occludin and positively correlating with ZO-1, but not JAM-A, Cingulin or ESAM (Supplemental figure 7B).

Immunohistochemistry analysis of the ear dermis similarly showed a downregulation of ZO-1 and an upregulation of VE-Cadherin in *Cldn5* iECKO mice (Figure 5C and D). Analysis of Occludin in these samples was precluded by its lack of expression in this vascular bed (Figure 1E). VE-Cadherin and ZO-1 expression was further investigated to determine in which vessel types, according to αSMA expression, their change in expression is occurring. ZO-1 expression was found to be significantly decreased specifically in venules whilst VE-Cadherin was equally upregulated in both capillaries and venules, but not in arterioles (Figure 5Eii).

Claudin5 expression thus regulates the expression and localisation of other EC junction components. Consequently, whilst Claudin5 loss results in no overt changes in junction structure it alters the regulatable and dynamic nature of the remaining junctional complex leading to changes in barrier stability and stringency.

## Discussion

The EC barrier consists of a variety of transmembrane cell adhesion molecules, allowing the formation of blood vessels with junctions of highly variable composition and strictness. However, the physiological consequences of this variability nor the mechanisms contributing to this variability are unclear. In this study we investigated the heterogeneous nature of the BEC barrier and uncover its composition and integrity in different organs. We determined that non-CNS, continuous endothelia broadly share a similar complement of AJ and TJ genes. Still, subtle variation in their relative distribution between different vessel subsets are often evident, as is variability in their barrier integrity and response to stimulation. Differential gene expression analysis revealed differences in gene expression between vessel subsets within tissues, which is provided as a resource along with expression values of genes typically associated with junction regulation and integrity (Supplemental figures 3 and 5 and Supplemental data 1 and 2). Generally, transmembrane TJ genes were found to be expressed to a greater degree in the arterial aspect of the microvasculature. In particular, *Cldn5* stands out in this analysis as it exhibited a large decrease in expression from arterioles through capillaries to venules in all tissues analysed, including in the human dermis. Previously we have shown that, in the ear skin, Claudin5 expression inversely correlates with susceptibility to VEGF-A-induced leakage, with capillaries possessing a salt-and-pepper mixture of Claudin5-positive and -negative cells being responsive to leakage stimulation (Honkura et al., 2018). This correlation however does not extend to other vascular beds such as the back skin and trachea, in which the leakage permissive vasculature was shifted away from the salt-and-pepper capillary region towards the venous side. To fully address why Claudin5-negative capillaries in the trachea might be resistant to leakage requires more extensive insights into the junctional organisation in these organs. Of note, whilst tracheal capillaries might be resistant to leakage, the trachea shows a much larger leakage response than the ear skin. The back skin similarly showed a larger leakage response than the ear skin and a more distinct separation between diminishing Claudin5 expression and leakage susceptibility. These data reveal the highly heterogeneous nature of barrier integrity and patterning across different tissues.

Differential regulation of EC barrier integrity in tissue-specific vasculatures was also observed following removal of Claudin5 in adult mice. We find that outside of the CNS, the largest influence of Claudin5 is in the ear and back skin vasculatures among the tissues analysed, although with varying impact on the passage of different tracers. In the ear skin Claudin5 protects barrier integrity against molecules larger than 2000 kDa dextran (approx. 54 nm diameter (Armstrong et al., 2004)), but not in the back skin, where Claudin5 only limits the passage of larger 200 nm particles. This is in stark contrast to the CNS vasculature where Claudin5 limits the passage of molecules as small as 443 Da (Nitta et al., 2003).

EC barrier integrity in skeletal muscle only showed minor change following the loss of Claudin5. This observation might be explained by the expression patterning of *Cldn5* in skeletal muscle, which declines faster along the arteriovenous axis in skeletal muscle than tissue such as the ear skin, leading to a 15-fold change in arterial/venous *Cldn5* expression in skeletal muscle versus a 3-fold change in ear skin (Figure 1F). The trachea vasculature similarly shows a large decrease in *Cldn5* expression in post-arterial BECs. Skeletal muscle BECs do however express other EC junction genes, such as *Ocln, Nectin2* and *Nectin3*, which are not evident in ear skin and tracheal BECs and might influence junction integrity (Figure 1E and Supplemental figure 3). It is currently uncertain what role, if any, these proteins have in EC junction strength. Studies have demonstrated a barrier protective role for Occludin, nectin-2 and nectin-3 *in vitro* (Martin et al., 2013, Son et al., 2016, Murakami et al., 2009). Further study is required to establish whether these actively participate in EC barrier integrity *in vivo* (Saitou et al., 2000).

In keeping with previous observations in the brain, loss of Claudin5 did not alter BEC junction structure in the ear dermis (Nitta et al., 2003). Smilarly, loss of VE-Cadherin in the lungs enhances barrier permeability without causing any structural defects (Duong et al., 2020). Interestingly, concomitant loss of ESAM along with VE-Cadherin produces more overt changes in junction structure and enhances loss of barrier integrity, highlighting the redundant organisation of the EC barrier (Duong et al., 2020). Interpretation of models that manipulate the EC barrier are often complicated by compensatory effects and redundancy. For example, VE-cadherin has been found to be a major regulator of EC gene expression (Morini et al., 2018). Interestingly, assembly of VE-Cadherin junctions upregulates Claudin5 expression, whilst its loss leads to compensatory upregulation of N-Cadherin (Taddei et al., 2008, Giampietro et al., 2012). JAM-A levels have similarly been found to positively regulate Claudin5 expression (Kakogiannos et al., 2020). We find here *in vivo* that Claudin5 can reciprocally alter expression of VE-Cadherin, as well as Occludin and ZO-1. The upregulated VE-Cadherin levels may explain the lack of structural defects in *Cldn5* iECKO mice. In contrast to Claudin5, VE-Cadherin is known to be highly dynamic and its localisation at junctions is regulated by numerous cytokines and growth factors that affect its phosphorylation status (Smith et al., 2020, Orsenigo et al., 2012, Eliceiri et al., 1999). Replacement of Claudin5 with VE-Cadherin is therefore likely to produce junctions that are less stable and subject to higher turnover in a stimulatory environment.

Collectively, the data in this study highlights the variable regulation and integrity of the EC barrier in distinct vascular beds and demonstrates the ability of Claudin5 to assemble junctions with different size-selective properties in different tissues. The mechanisms underlying such organotypic barrier integrity and differential patterning of junctional components are currently poorly understood. Pericytes are known to be essential for maintenance of the tight BBB (Armulik et al., 2010). Outside of the CNS pericyte coverage is comparatively low, whether coverage differs between non-CNS vascular beds however is unknown. Barrier integrity may also be regulated by basement membrane components such as laminin α5, which enhances VE-Cadherin stability at cellcell junctions (Richards et al., 2021, Song et al., 2017). Similar to pericytes, basement membrane coverage differs between different vascular beds, as well as between vessel subtypes (Richards et al., 2021, Di Russo et al., 2017). Whether basement membrane components significantly define EC junction composition however is currently unknown. A better understanding of basement membrane, as well as pericyte, coverage however would provide potential new means of manipulating EC biology for therapeutic benefit.

In conclusion, this study provides an in-depth characterisation and comparison of the EC barrier in numerous organs. Furthermore, we demonstrate that Claudin5 plays an organotypic role EC barrier integrity. In accordance with its role in forming the tight BBB and BRB in the CNS, we find that Claudin5 influences vascular permeability in the skin, albeit in a size-selective manner that is dramatically different from the CNS. The impact of Claudin5 in organ-specific vascular permeability was demonstrated by the direct correlation between its expression and high barrier integrity in some, but not all vascular beds. Variability in the pattering of the EC barrier between vascular beds and vessel subtypes has remained poorly defined, as is the redundancy, interdependency and crosstalk between different junction components. Understanding the role of different EC components in various vascular beds will allow us to better appreciate organ-specific defects in vascular permeability and how they may be therapeutically targeted.

## Acknowledgements

The authors acknowledge the Biocenter Oulu Electron Microscopy Core Facility supported by Biocenter Finland and the University of Oulu for their specific scientific expertise and research infrastructure services and the equipment and expert advice supplied by the BioVis imaging and flow cytometry core facility (Uppsala University). This study was supported by the Swedish Research Council (2020-01349), the Knut and Alice Wallenberg foundation (KAW 2020.0057 and KAW 2019.0276), Fondation Leducq Transatlantic Network of Excellence Grant in Neurovascular Disease (17 CVD 03) and the Swedish Cancer foundation 19 0119 Pj and 19 0118 Us to L.C.-W. K.K. and M.G. were supported by Cancerfonden (20 1086 Pj), KK is supported by Wallenberg Academy Fellowship (2017.0144), Ragnar Söderbergs Fellowship (M13/17). E.S. was supported by Svenska Sällskapet för Medicinsk Forskning (SSMF). E.N. was supported by the Gustaf Adolf Johansson’s foundation. S.N. is supported by Åke Wibergs foundation (M21-0109). M.R. was supported by SSMF (201912) and an EMBO long-term fellowship (ALTF 923-2016).

## Author Contributions

Conceptualization, M.R., S.N. and L.C.-W.; methodology, M.R., E.N. and S.N.; investigation, M.R., E.N., S.P., P.M., M.K., M.G., E.S. and S.N.; writing, M.R., S.N. and L.C.-W.; funding acquisition; M.R. and L.C.-W.; resources, C.B., L.E., K.K. and L.C.-W.

## Supplemental figure legends

**Supplemental figure 1.**
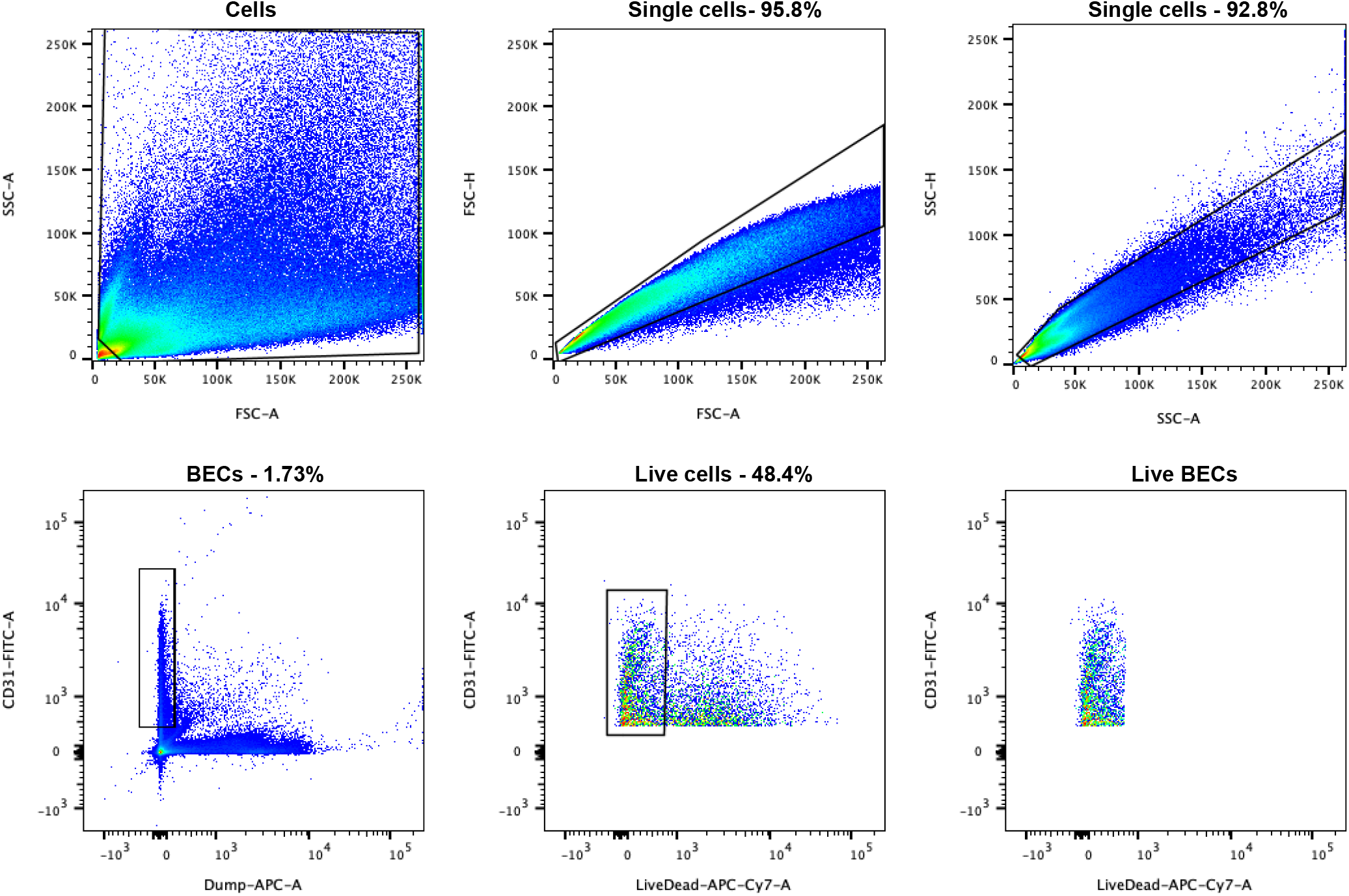
Gating strategy for the FACS isolation of single BECs from the mouse ear skin.

**Supplemental figure 2.**
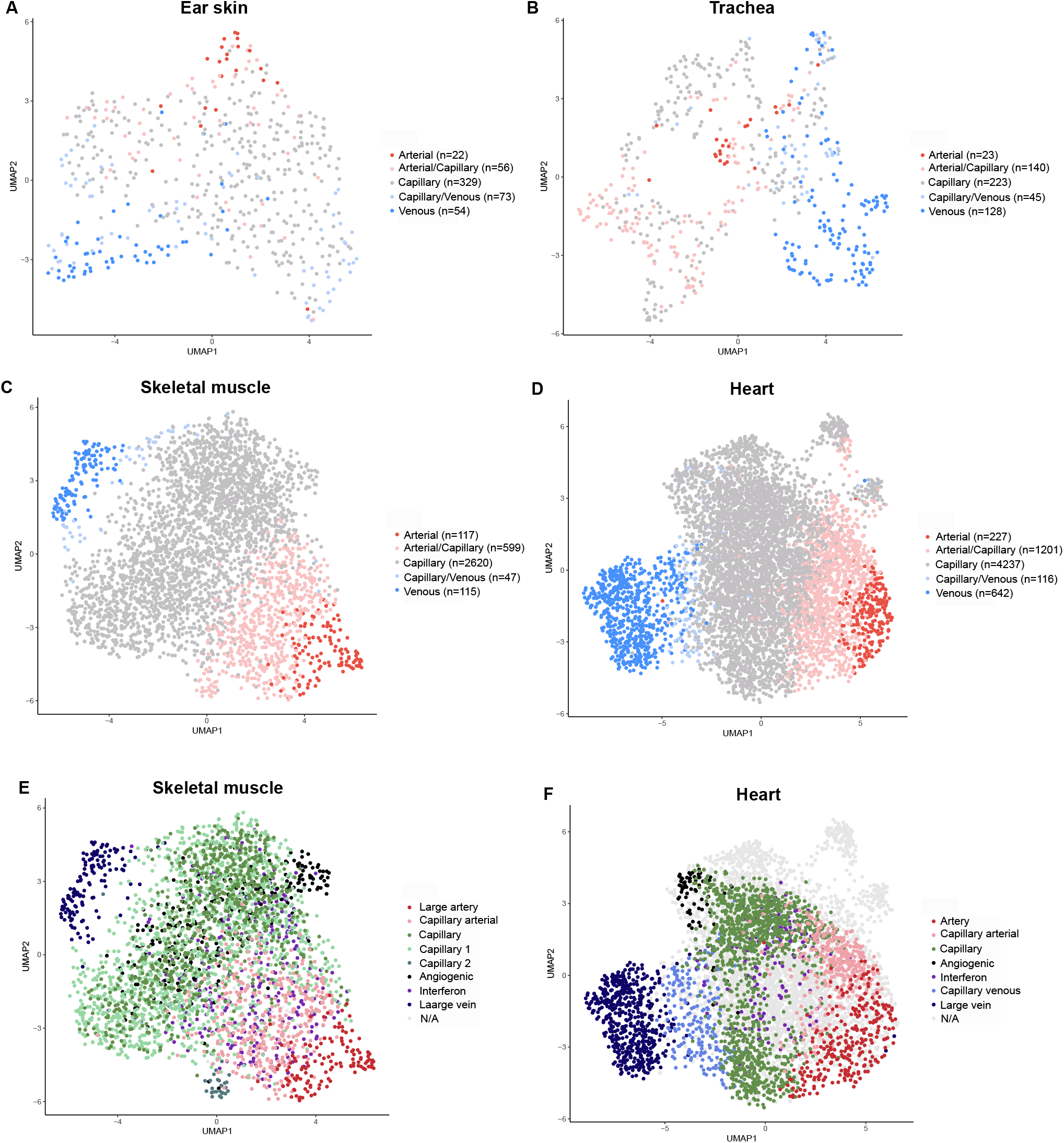
**A**. Ear skin, **B**. trachea, **C**. skeletal muscle and **D**. heart mouse BECs coloured by subset as defined by analysis of integrated data. The number of cells in each subtype are specified in each legend. **E**. Skeletal muscle and **F**. heart mouse BECs coloured by clusters identified in the original publication of the data (Kalucka et al., 2020).

**Supplemental figure 3.**
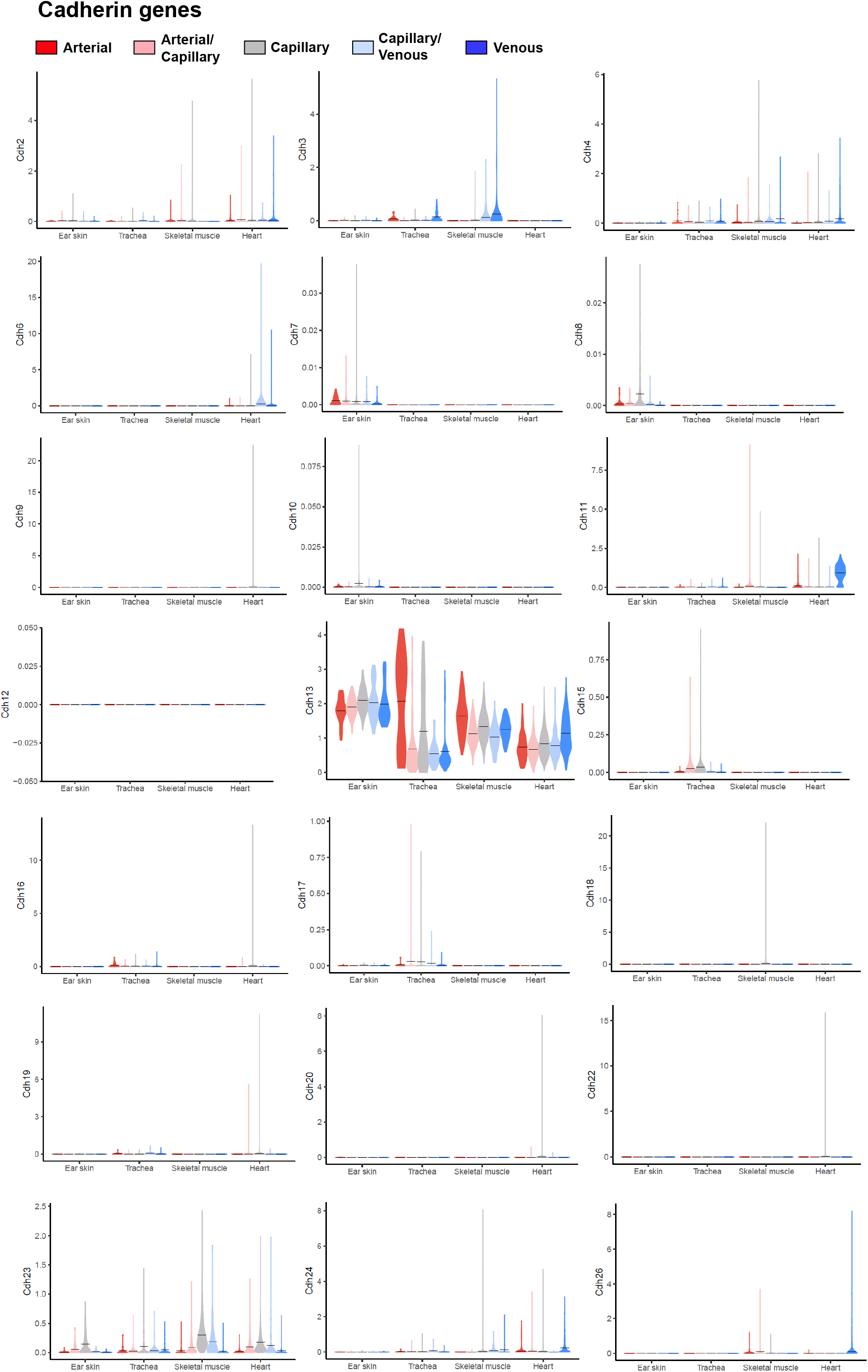
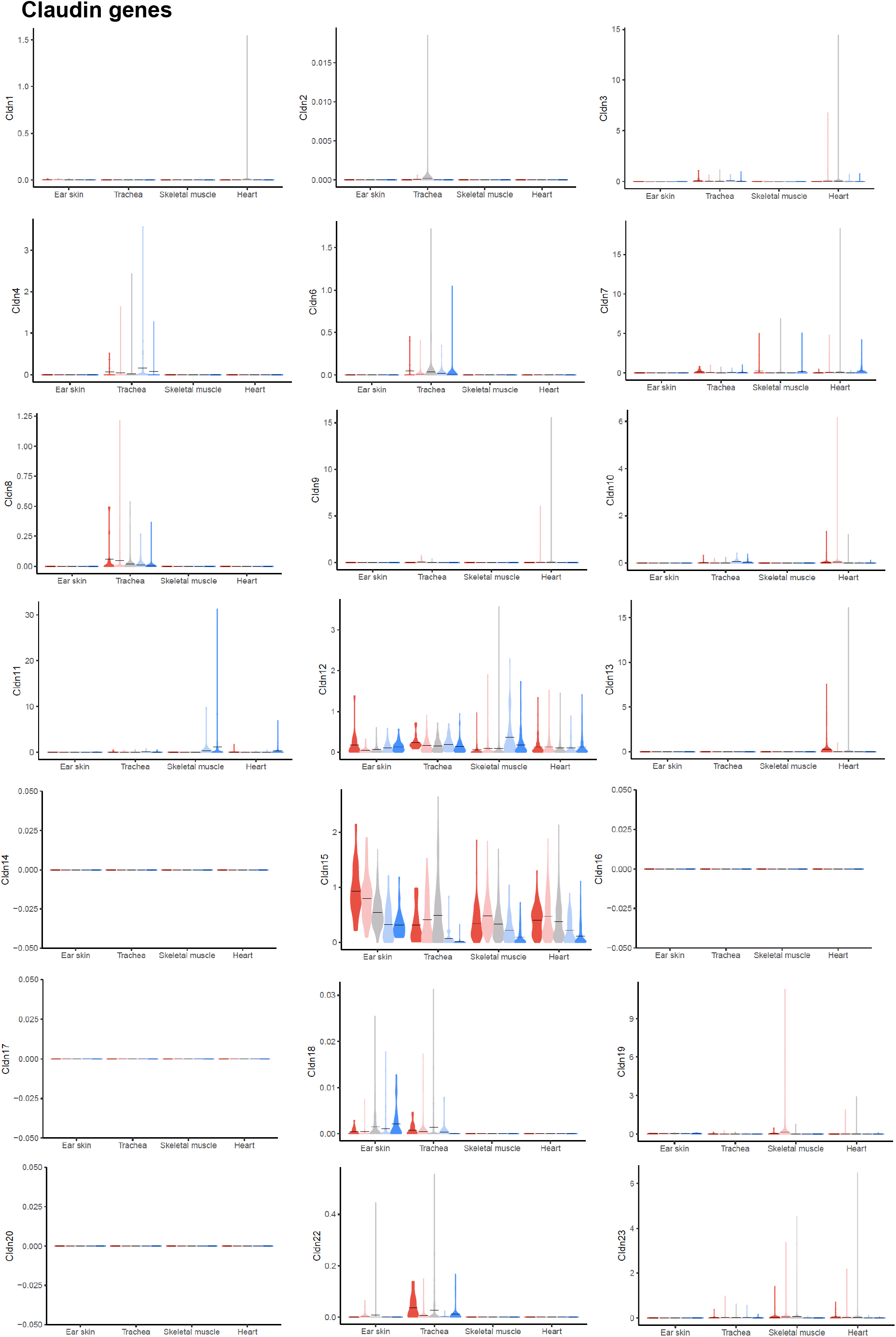
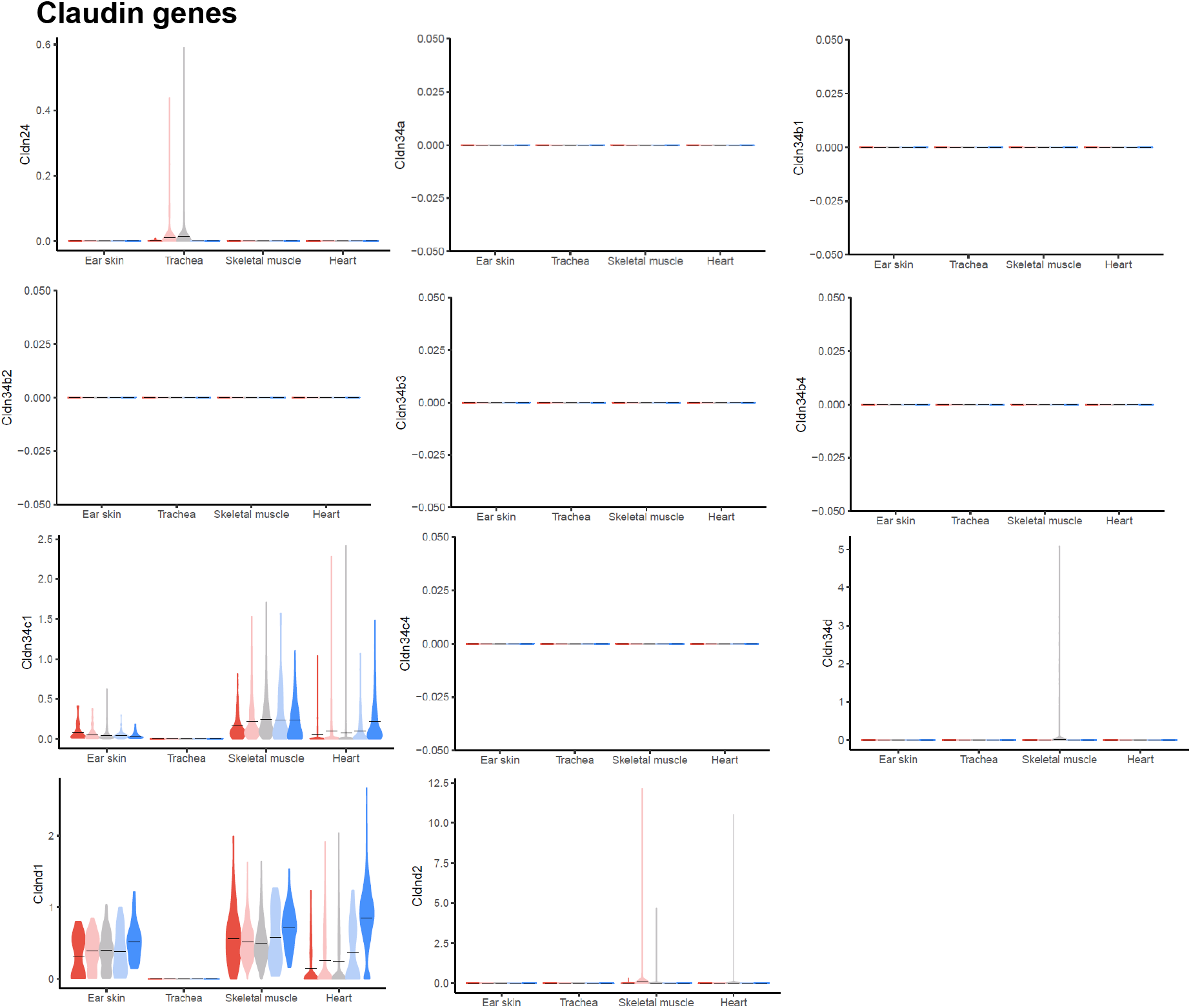
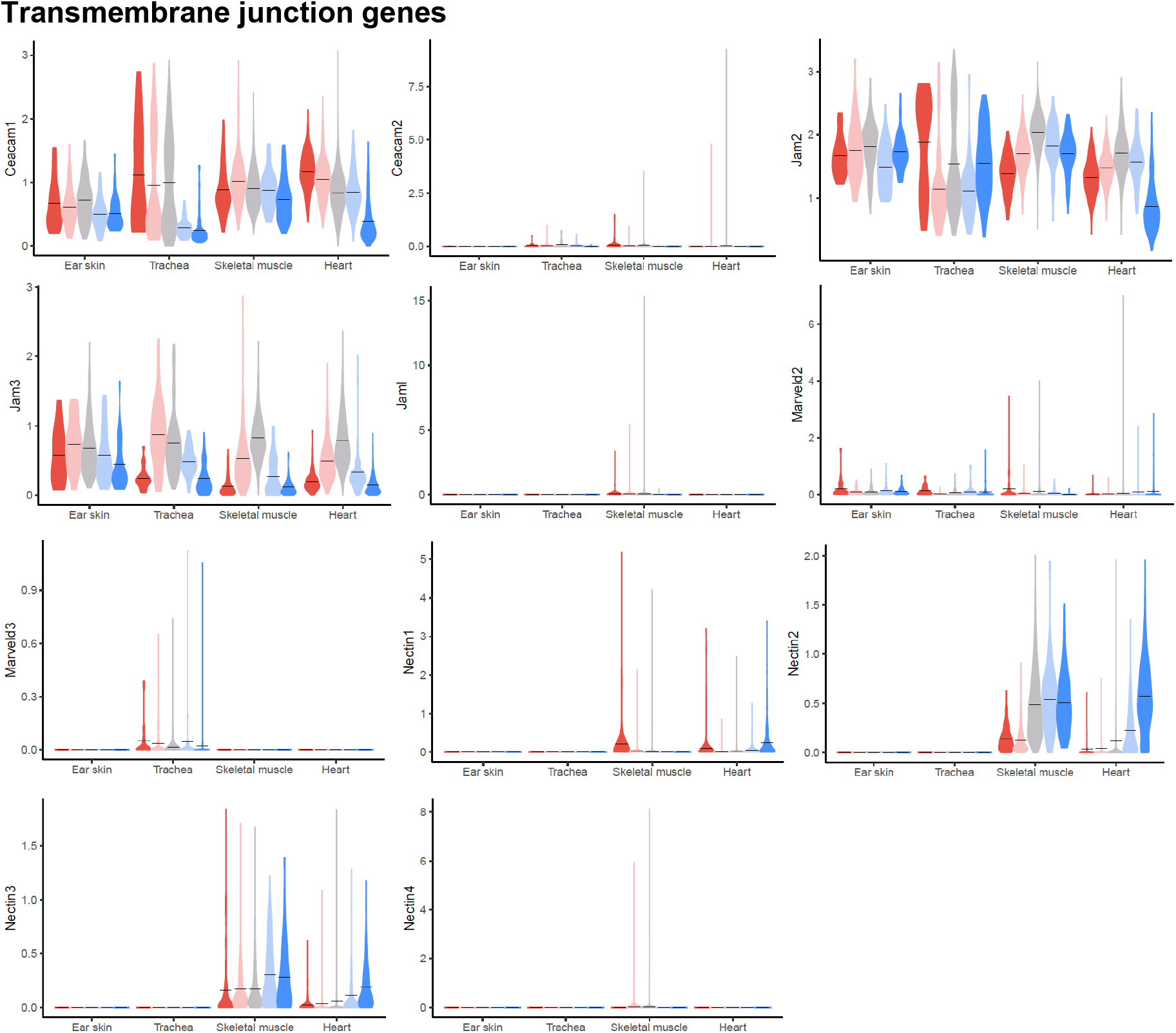
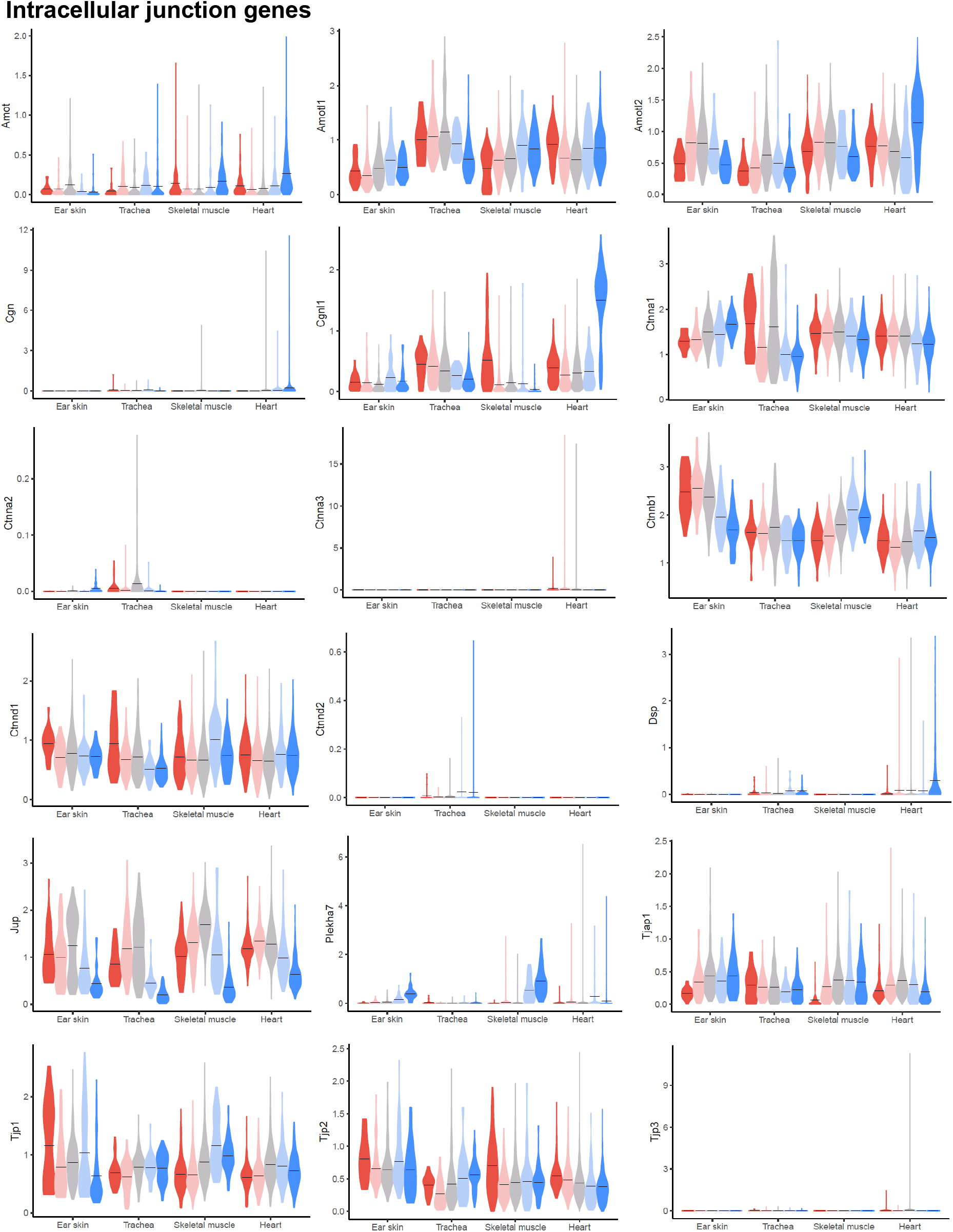
Violin plots of gene expression for mouse endothelial junctional components. Gene expression was normalized to account for differences in sample library size and imputed to account for dropouts in the data as described in Methods.

**Supplemental figure 4.**
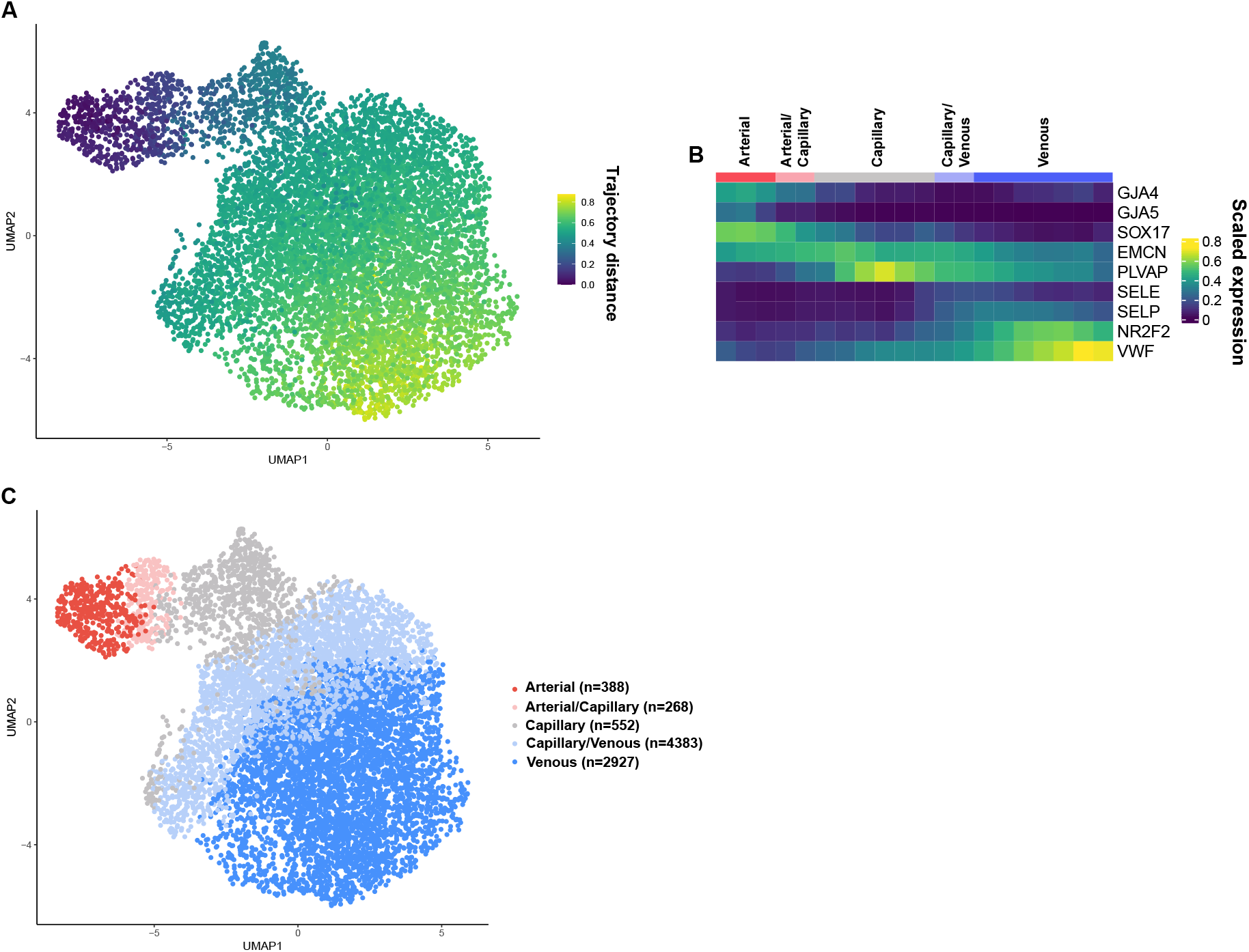
**A**. Uniform manifold approximation and projection (UMAP) of human dermal BECs showing the distance of an isolated trajectory calculated with tSpace. **B**. Equidistant binning of the trajectory shown in A. with supervised annotation of the bins as vessel subsets. **C**. UMAP coloured by the vessel subsets specified in B.

**Supplemental figure 5.**
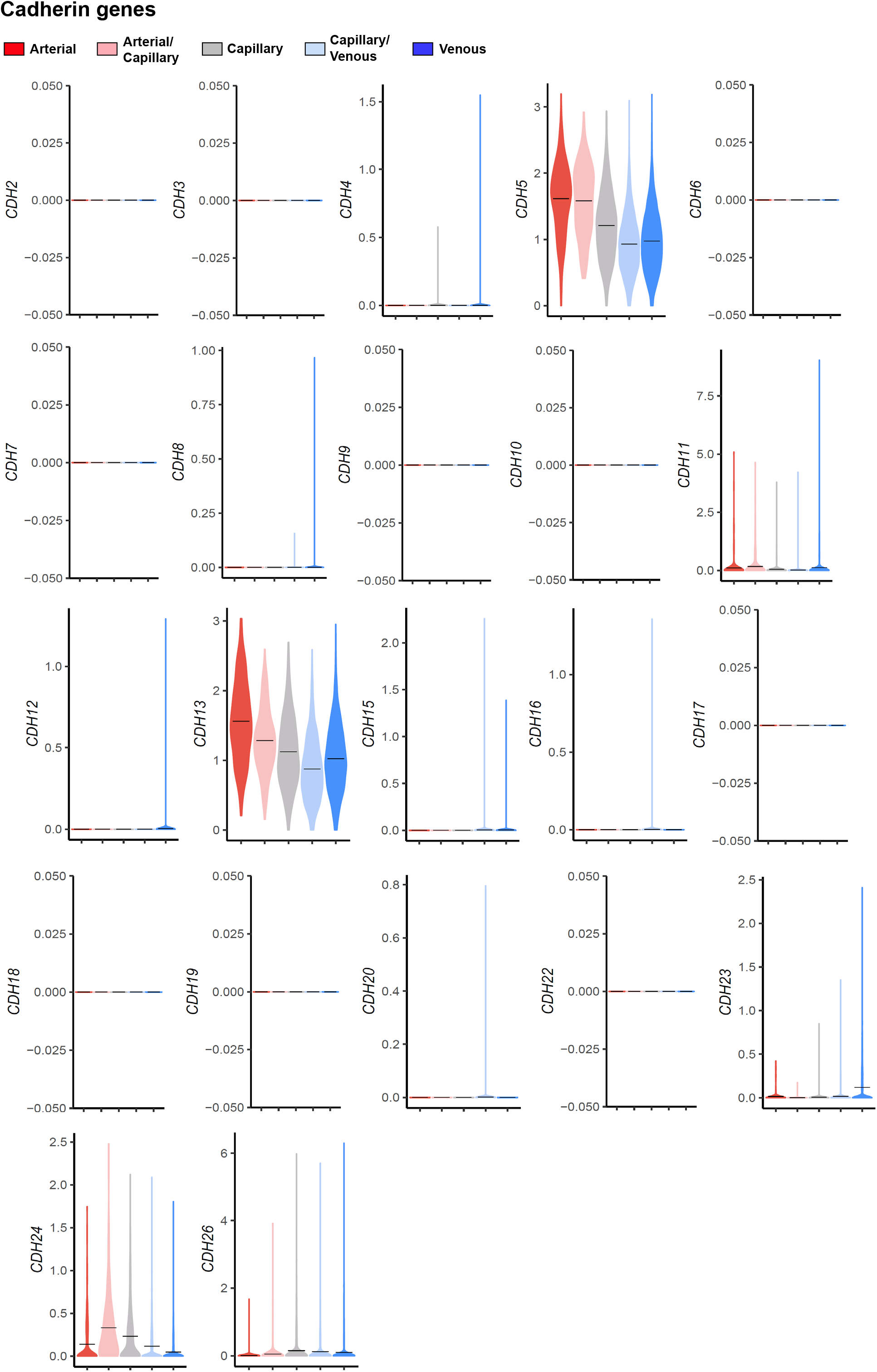
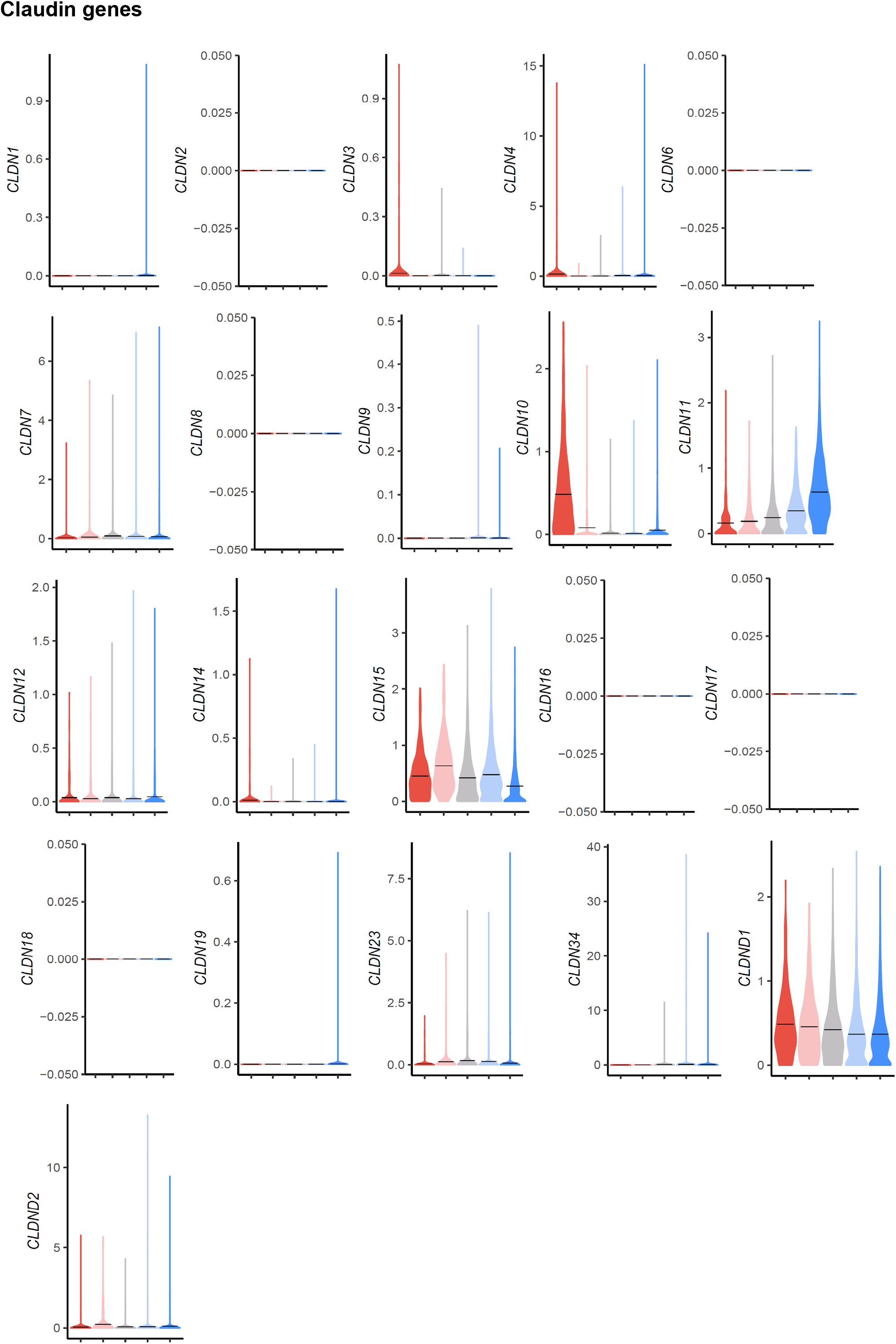
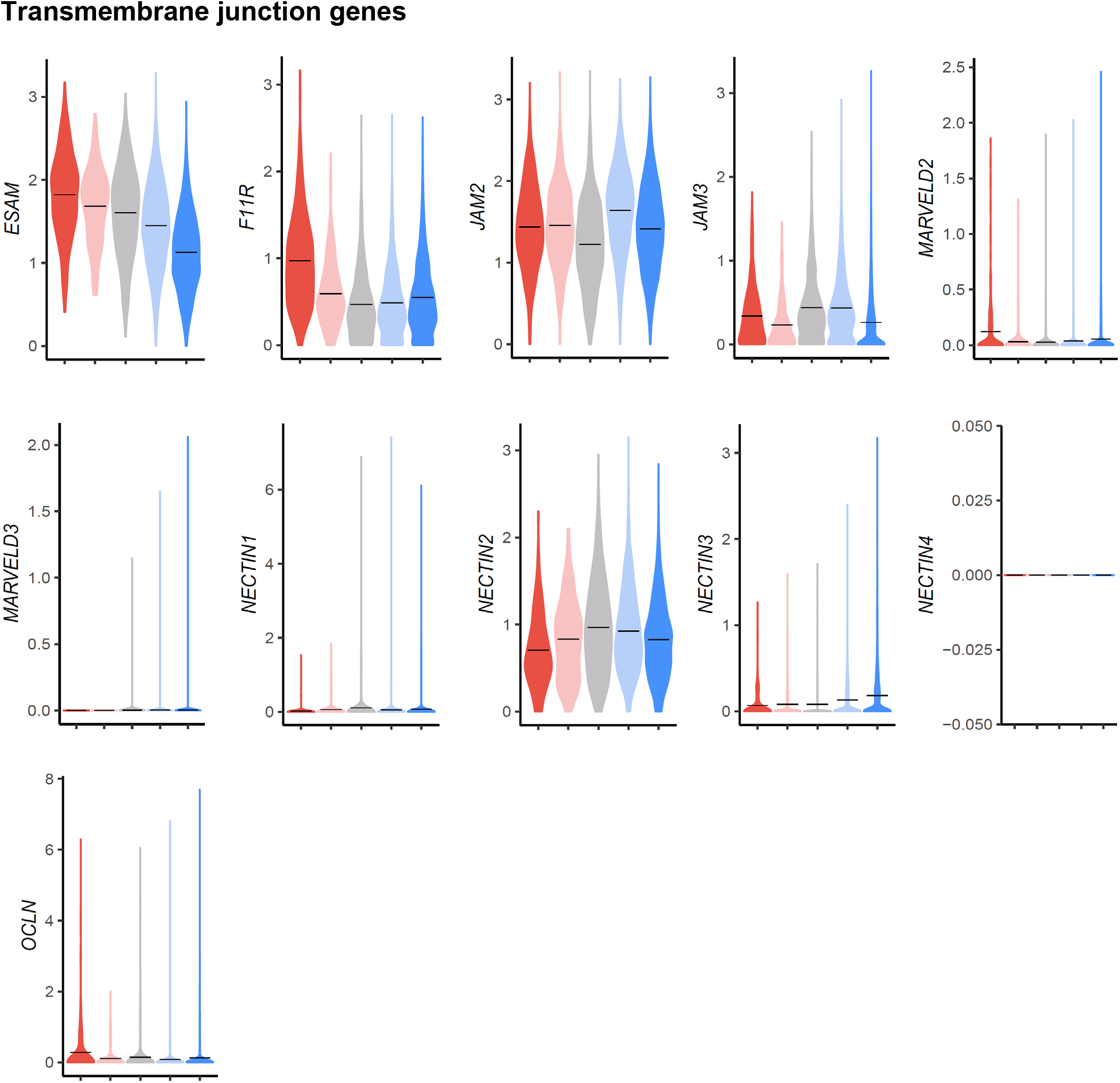
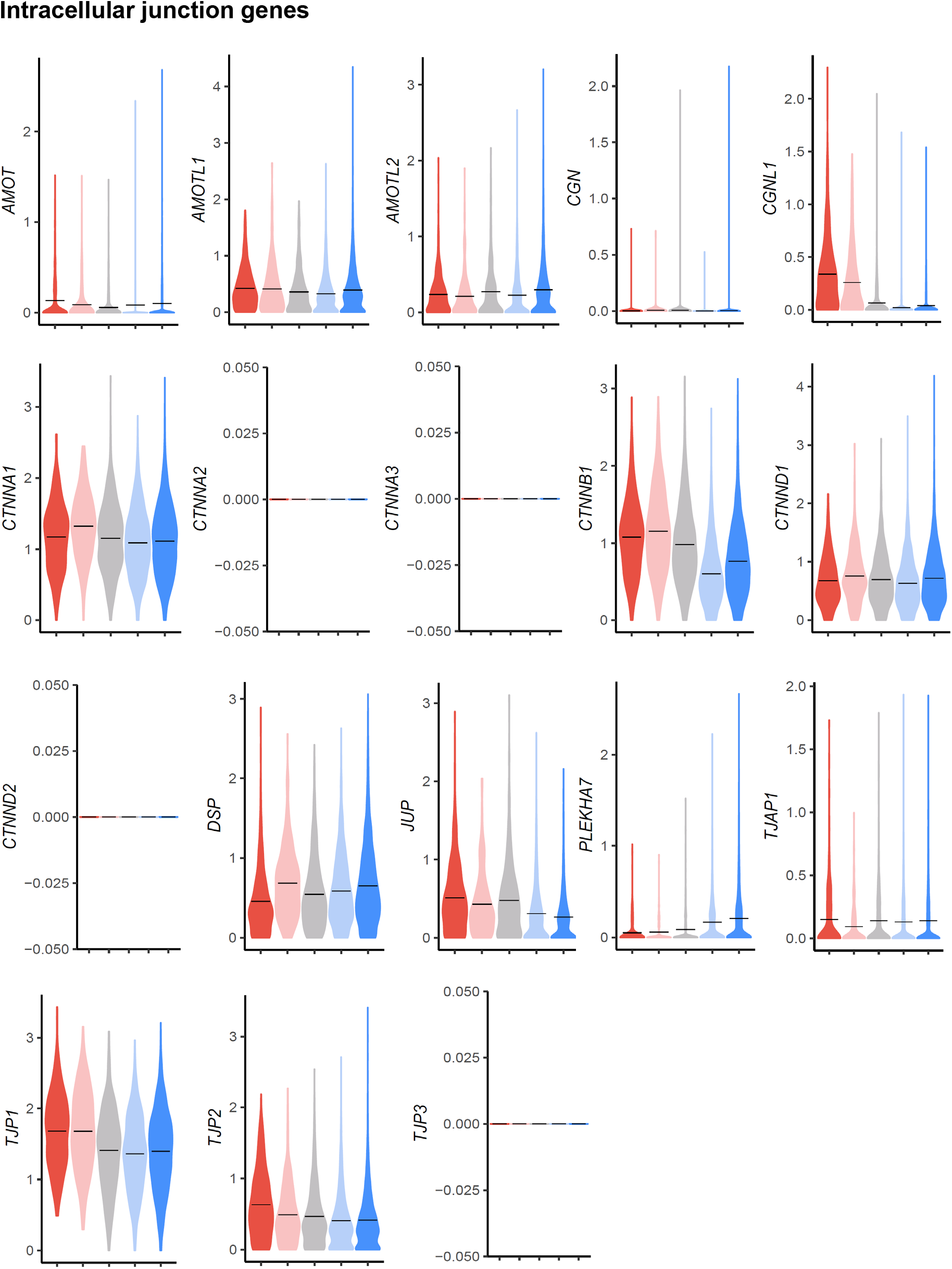
Violin plots of gene expression for human dermal endothelial junctional components. Gene expression was normalized to account for differences in sample library size been imputed to account for dropouts in the data as described in Methods.

**Supplemental figure 6.**
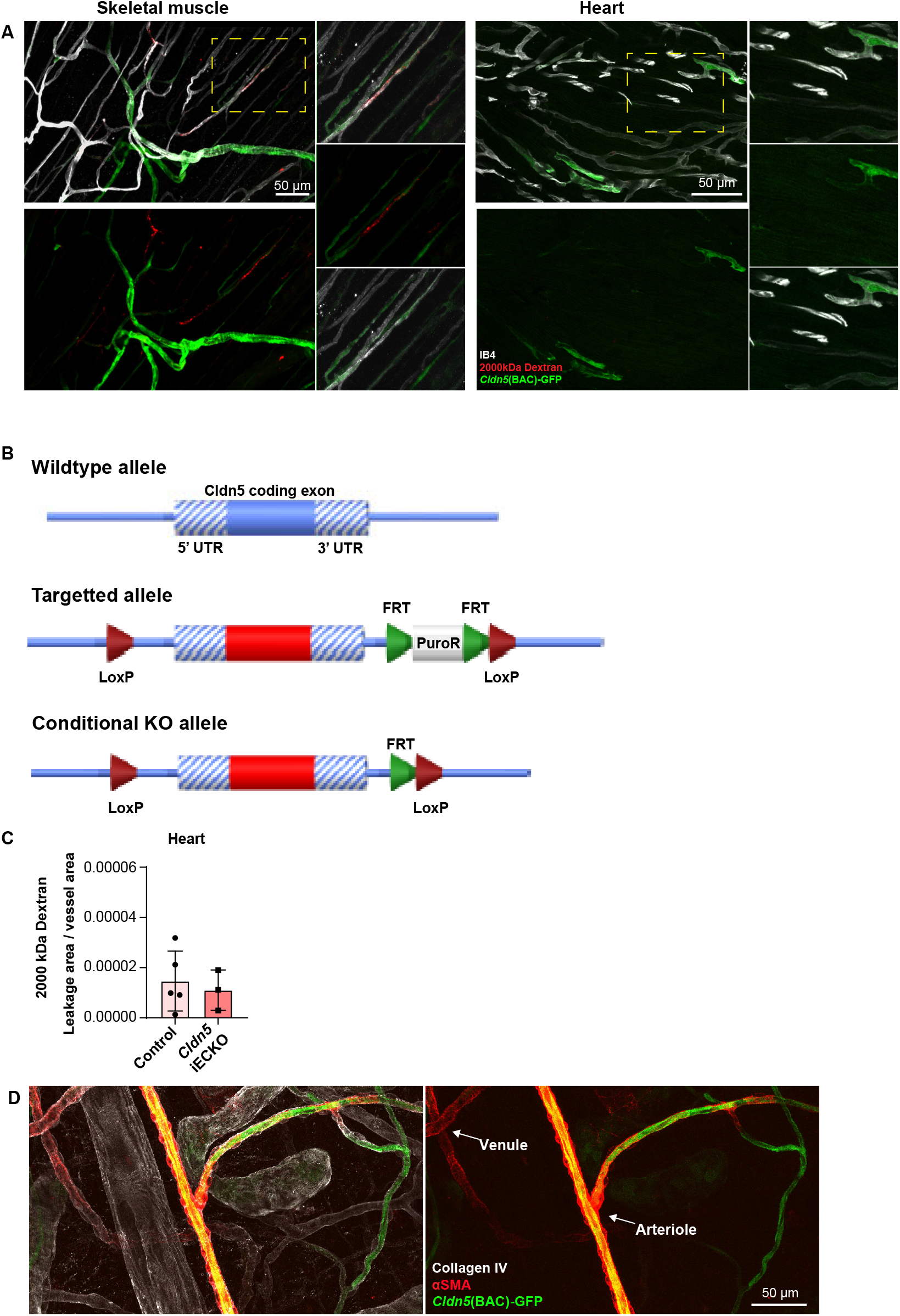
**A.** Leakage and *Cldn5*(BAC)-GFP expression patterning in skeletal muscle and heart in response to the systemic administration of histamine. Magnified images of dashed boxes are shown to the right of the main image. **B.** Schematic diagram showing the targeting strategy of *Cldn5* floxed mice. **C.** Quantification of histamine-induced leakage of 2000 kDa dextran from the heart vasculature in control and *Cldn5* iECKO mice. n ≥ 3 mice, 3 or more fields of view/mouse. **D.** Image showing the expression of *Cldn5*(BAC)-GFP and αSMA in the mouse ear dermis.

**Supplemental figure 7.**
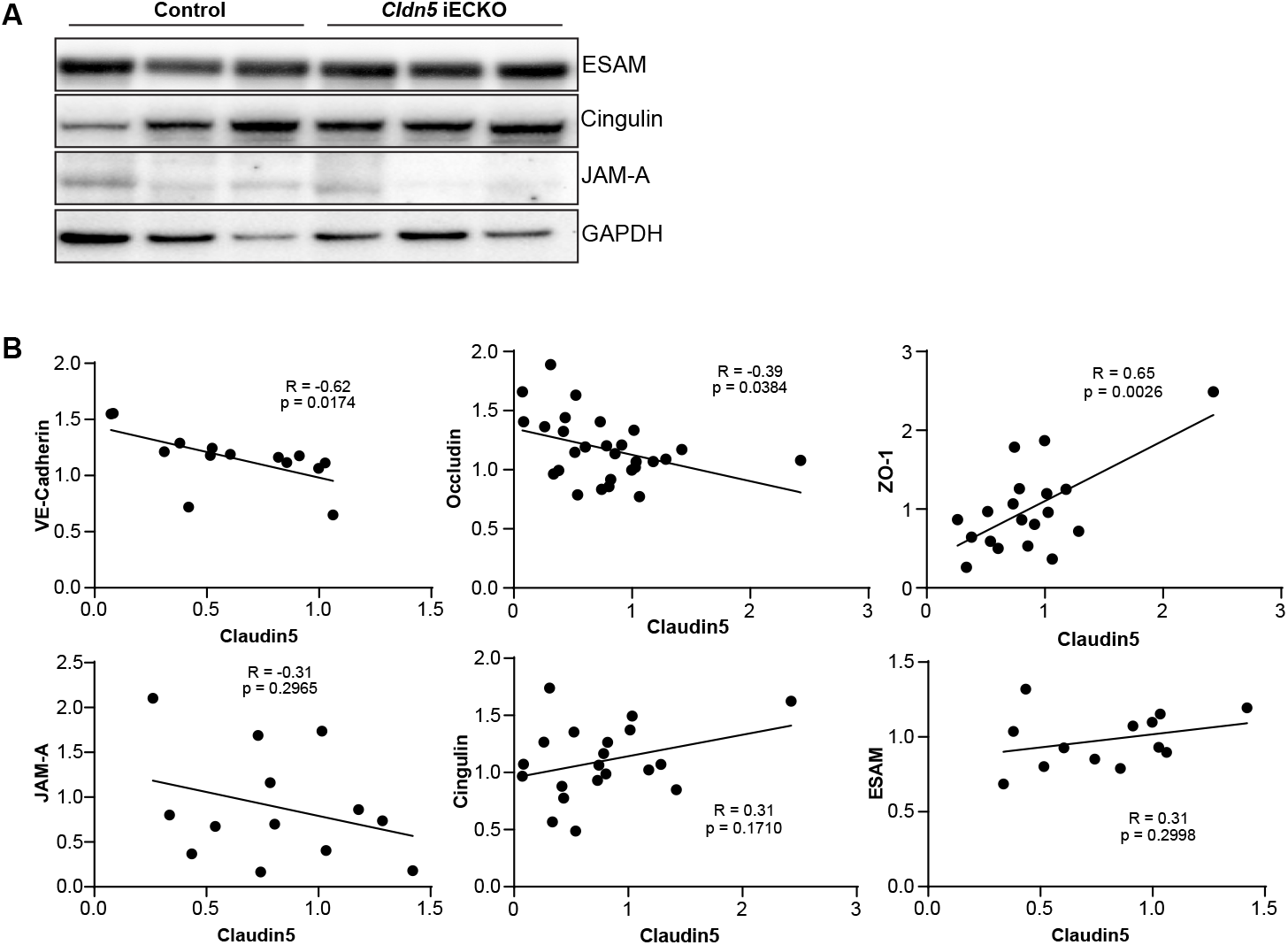
**A.** Representative western blots of ESAM, Cingulin and JAM-A expression in control and *Cldn5* iECKO mouse lung lysates. **B.** Scatter graphs showing the correlation between Claudin5 expression levels and other EC junction proteins in mouse lung lysates. Both control and *Cldn5* iECKO mice are represented. Line of best fit following linear regression analysis is shown.

<u>Supplemental data 1</u>

Spreadsheets detailing the results of the differential gene expression analysis conducted between mouse BEC subtypes in ear skin (1.1), trachea (1.2), skeletal muscle (1.3) and heart (1.4).

<u>Supplemental data 2</u>

Spreadsheet detailing the results of the differential gene expression analysis conducted between human dermal BEC subtypes.

<u>Movie 1: Histamine-mediated leakage in *Cldn5*(BAC)-GFP mice</u>

Extravasation of circulating 2000 kDa TRITC Dextran (pseudocolour) after intradermal injection of histamine in the ear dermis of *Cldn5*(BAC)-GFP mice. The first 30 frames show a still at t=0 to show *Cldn5*(BAC)-GFP expression (green) overlayed with 2000 kDa Dextran (grey).

<u>Movie 2: Histamine-mediated leakage in control and *Cldn5* iECKO mice</u>

Extravasation of circulating 2000 kDa FITC Dextran (pseudocolour) in control (left) and *Cldn5* iECKO (right) mice after intradermal injection of histamine in the ear dermis.

## Methods

### Animals

For generation of a mouse ear dermal scRNAseq dataset wild-type female C57BL/6J mice aged 8-18 weeks were used. *Cldn5*^flox/flox^ were generated by Taconic by flanking the sole exon with *loxP* sites, introduced by homologous recombination on the genetic C57Bl/6 black background (Supplemental figure 7). This strain was crossed with Cdh5(PAC)-CreER^T2^ mice (kind gift from Dr. Ralf Adam, Max-Planck Institute Münster) to generate endothelial specific knockout of *Cldn5. Cldn5*(BAC)-GFP mice have been described (Honkura et al., 2018, Lavina et al., 2018). To monitor Cre recombinase activity with YFP expression the fluorescent reporter mouse line B6.129X1-Gt(ROSA)26Sor^tm1(EYFP)Cos^/J (Stock Number 006148, The Jackson Laboratory) was introduced. Both males and females, aged 8-18 weeks, were included in experiments. *In vivo* animal experiments were carried out in accordance with the ethical permit provided by the Committee on the Ethics of Animal Experiments of the University of Uppsala (permit 6789/18). Mice were maintained in ventilated cages with group housing (2-5 per cage). Each experiment was conducted on tissue from at least three age-matched animals representing individual biological repeats. Sample size (number of acquired images / movies and number of mice) were chosen to ensure reproducibility and allow stringent statistical analysis. To induce Cre recombinase-mediated gene recombination tamoxifen (SigmaAldrich, T5648) was injected intraperitoneally (1mg/day) for 5 consecutive days. The mice were allowed to rest for 5 days before experiments were conducted. For topical tamoxifen induction 50 μg of 4-hydroxytamoxifen (SigmaAldrich, H7904) in acetone was applied to each side of the mouse ear for 3 consecutive days. Mice were allowed to rest for 14 days before experiments were conducted.

### Intravital vascular leakage assay

Intravital imaging of the mouse ear with intradermal injection has been described previously (Honkura et al., 2018). Briefly, following systemic administration of 2000kDa FITC (SigmaAldrich, FD2000S) or TRITC Dextran (ThermoFischer Scientific, D7139) by tail-vein injection, mice were anaesthetised by intraperitoneal injection of Ketamine-Xylazine (120 mg/kg Ketamine, 10 mg/kg Xylazine) to a surgical level and the ear secured to a solid support. Mice were maintained at a body temperature of 37°C for the entire experiment, maximum 90 minutes. Time-lapse imaging was performed using single-photon microscopy (Leica SP8). For intradermal EC stimulation, a volume of approximately 0.1 μl histamine (SigmaAldrich, H7125), concentration 10 ng/μl, was injected using a sub-micrometer capillary needle. 10 kDa TRITC Dextran (ThermoFischer Scientific, D1817) was used as a tracer. Leakage sites were identified in time-lapse imaging as defined sites of concentrated dextran in the extravascular space.

### Permeability analysis

To assess EC permeability all tissues were cleaned of excess cartilage, fat and connective tissue. Skeletal muscle was assessed using the tibialis anterior.

To analyse baseline permeability mixtures of dextran (Tdb labs; 4kDa, TD4, 10kDa, FD10, 70kDa, TD70) (10mg / 25g body weight) were injected systemically through the tail vein. Four hours later mice were anaesthetised with Ketamine/Xylazine before intracardiac perfusion with Dulbecco’s phosphate buffered saline (DPBS). Tissues were dissected, washed in DPBS and incubated in formamide for 48 hours at 37°C. Dextran fluorescence was then measured using a spectrophotometer and normalised to tissue weight. To assess organotypic induced leakage histamine (0.25 mg / 25g body weight) was mixed with 2000kDa dextran (SigmaAldrich, FD2000S) before systemic injection through the tail-vein. Dextran extravasation was assessed as above ten minutes later.

To assess histamine induced leakage microscopically mixtures of 2000kDa lysine fixable dextran (FITC, Tdb labs; TRITC, ThermoFischer Scientific, D7139) (10mg / 25g body weight) or fluorescent microspheres (ThermoFischer Scientific, B200) (75 μl; 1% solids / 25g body weight) with histamine (0.25 mg / 25g body weight) were injected systemically through the tail vein. Ten minutes later mice were anaesthetised with Ketamine/Xylazine before intracardiac perfusion with DPBS followed by 1% paraformaldehyde. Tissues were then immersed in 1% paraformaldehyde for 2 hours before proceeding with immunohistochemistry. For leakage quantification at least 3 large tile scan areas (≥ 1 mm^2^) were captured for each mouse.

### Immunohistochemistry

Tissues were fixed through intracardiac perfusion with 1% paraformaldehyde (PFA) followed by immersion in 1% PFA for 2 hours at room temperature. Tissues were blocked overnight at 4°C in Trisbuffered saline (TBS), 5% (w/v) Bovine Serum Albumin (BSA), 0.2% Triton X-100. Samples were incubated overnight with primary antibody in blocking reagent, followed by washing in TBS, 0.2% Triton X-100 and incubation with appropriate secondary antibody for 2 hours at room temperature in blocking buffer before washing and mounting in fluorescent mounting medium (DAKO). Images were acquired using a Leica SP8 confocal microscope.

Whole mount preparation of the ear skin, back skin and trachea were made following the removal of excess cartilage, fat and connective tissue. Assessments of skeletal muscle (tibialis anterior) and heart tissues were made using 100μm longitudinal vibratome sections.

Commercial antibodies used were: rat anti-CD31 (BD Biosciences, 553370), goat anti-CD31 (R&D Systems, AF3628), goat anti-VE-Cadherin (R&D Systems, AF1002), chicken anti-GFP (Abcam, ab13970) (1:400), rabbit anti-Claudin5 (ThermoFischer Scientific, 341600), rabbit anti-ZO-1 (ThermoFischer Scientific, 617300), goat anti-Collagen IV (Merck Millipore, AB789), mouse anti-αSMA FITC (Sigma Aldrich, F3777), mouse anti-αSMA Cy3 (Sigma Aldrich, C6198), *Griffonia simplicifolia* Isolectin GS-IB4 (Molecular Probes, I32450) (1:400). Secondary antibodies against rat (ThermoFischer Scientific; Alexa 488, A21208 and Alexa 594, A21209), rabbit (ThermoFischer Scientific; Alexa 488, A21206 and Alexa 568, A10042), goat (ImmunoResearch Laboratories, Alexa 647, 705-605-147) and chicken (ImmunoResearch Laboratories, Alexa 488, 703-545-155) were used. Primary and secondary antibodies were prepared at a dilution of 1:100 and 1:400 respectively unless otherwise stated.

### *In situ* RNA hybridisation

Detection of *Cldn5* mRNA in the ear dermis was performed using RNAscope Fluorescent Multiplex Assay (ACD Bio, 322340 and 320851) according to the manufacturer’s instructions (www.acdbio.com). Briefly, 10μm transverse cryosections of the ear dermis were fixed in chilled 4% PFA for 15 minutes (4°C) then dehydrated by incubating in increasing concentrations of ethanol (50%, 75%, 100%, 100% for 5 minutes each). Following protease IV digest (30 minutes at RT), Cldn5 probe (ACD Bio, 491611-C2) was hybridized on the tissue sections for 2 hours at 40°C. 3-plex negative (ACD Bio, 320871) and positive (ACD Bio, 320811) controls were used to confirm signal specificity. Signal amplification and detection was conducted using reagents included in the Fluorescent Multiplex Reagent Kit. Following the RNAscope procedure, sections were immediately counterstained with Hoechst (Molecular Probes, H3570) (1:1000), *Griffonia simplicifolia* Isolectin GS-IB4 (Molecular Probes, I32450) and αSMA (Sigma Aldrich, F3777) (1:300) for 1 hour at RT to visualize nuclei, blood vessels and mural cells, respectively. Images were acquired using a Leica SP8 confocal microscope with at least 3 sections analysed / mouse. Fluorescent dots representing one mRNA molecule were quantified in *Cldn5*(BAC)-GFP-negative and -positive vessels before normalising to these vessel areas. Images were acquired using a Leica SP8 confocal microscope with at least 3 sections analysed / mouse. Fluorescent dots representing one mRNA molecule were quantified in *Cldn5*(BAC)-GFP-negative and -positive vessels before normalising to these vessel areas.

### Ear dermal single cell isolation

Following cervical dislocation, ears of wild-type C57BL/6J mice were collected and mechanically disrupted before enzymatic dissociation in 10 mg/ml Collagenase IV (Worthington, LS004183), 0.2% FBS and 0.2 mg/ml DNase I (Worthington, LS006333) in PBS at 37 °C for 20 minutes. Following removal of debris using a 50 μm filter, cells were resuspended in FACS buffer (0.5% FBS, 2mM EDTA in PBS) and incubated with an anti-CD16/32 antibody (ThermoFischer Scientific, 14-0161-85) on ice for 10 minutes. Subsequently, cells were incubated with antibodies against CD31-FITC (BD Biosciences, 553372), CD45-APC (BioLegend, 103112) and Lyve1-eFluor 660 (ThermoFischer Scientific, 50-0443-82) for 30 minutes before washing and addition of live/Dead near IR cell stain (ThermoFischer Scientific, L10119) to distinguish viable cells. CD31+/CD45-/Lyve1-cells were obtained by FACS following filtration through a 50 μm mesh using a BD FACSAria III (100 μm nozzle size, 20 psi sheet pressure) at the BioVis core facility at the Department of Immunology, Genetics and Pathology (IGP), Uppsala University. Cells were captured directly into 2.3 μl lysis buffer (0.2% Triton-X (Sigma, cat: T9284), 2U/μl RNase inhibitor (ClonTech, cat: 2313B), 2 mM dNTP’s (ThermoFisher Scientific, cat: R1122), 1 μM Smart-dT30VN (Sigma), ERCC 1:4 × 10^7^ dilution (Ambion, cat: 4456740) in 384-well plates prior to library preparation.

### Smart-seq2 library preparation and sequencing

Single-cell libraries were prepared as described previously (Picelli et al., 2014), with the following specifications: 0.0025 μl of a 1:40,000 diluted ERCC spike-in concentration stock and all cDNA was amplified with 22 PCR cycles before QC control with a Bioanalyzer (Agilent Biosystems). The libraries were sequenced on a HiSeq2500 at the National Genomics Infrastructure (NGI), Science for Life Laboratory, Sweden, with single 50-bp reads (dual indexing reads). All single-cell transcriptome data were generated at the Eukaryotic Single-cell Genomics facility at Science for Life Laboratory in Stockholm, Sweden.

### Data processing

Raw data for ear skin sequencing data was aligned to the mouse reference genome mm10 with tophat (v.2.1.1) (Kim et al., 2013), duplicated reads were filtered out using the samtools software (v.0.1.18), and gene counts were summarized using *featureCounts* function from the Subread package (v.1.4.6-p5) (Liao et al., 2014). Raw counts of trachea and heart BEC generated by the Tabula Muris Consortium (Tabula Muris, 2020) were obtained from https://s3.console.aws.amazon.com/s3/buckets/czb-tabula-muris-senis/. Both FACS (trachea) and droplet (trachea and muscle) processed data was used. Previously published heart and skeletal muscle data (Kalucka et al., 2020) was obtained from ArrayExpress (https://www.ebi.ac.uk/arrayexpress/), accession number E-MTAB-8077, and preprocessed with cellranger (v3.0.2) using the mm10 reference genome. Human dermal BEC data (Li et al., 2021) was obtained from the National Genomics Data Center (https://bigd.big.ac.cn/), accession number PRJCA002692, and preprocessed with cellranger (v5.0.1) using the GRCh38 reference genome.

BEC were defined by their expression of *Pecam1, Cdh5*, and *Kdr*. Non-BEC (defined by *Lyve1, Prox1, Ptprc, Pdgfrb*, or *Kcnj8* expression) and cells with a total count below 600, fewer than 500 or more than 8000 expressed genes, were removed from downstream analysis. A total of 534 (356+178) ear skin, 559 (356+203) trachea, 3498 (1118+2380) skeletal muscle, and 6423 (2923+3500) heart BEC were included in the analysis. 8518 BEC were identified and included in the human dermal dataset.

All processing steps were carried out with R (v.4.0.4; *Lost Library Book*) unless stated otherwise. Size factor-based normalization (McCarthy et al., 2017) and log transformation was carried out with scran (v.1.18.7) *quickCluster* and *computeSumFactors* followed by batchelor (v.1.6.3) *multiBatchNorm*, in order to adjust for differences in sequencing depth between samples; scuttle (v.1.0.4) *logNormCounts* was used when preparing the human dermal BEC data. Highly variable genes were identified with scran *modelGeneVar* using non-integrated data while blocking for sample specific bias. Imputation was done with the magic python package (v.0.1.1; k = 9, ka = 3, t = 1, 2, 4, or 6) in order to reduce the effects of dropouts, and integration of the datasets was done with batchelor *fastMNN* (k = 100) (van Dijk et al., 2018, Haghverdi et al., 2018). Trajectory inference was done using tSpace (v.0.1.0) and a total of 400 trajectories were calculated using imputed (t = 6) and integrated expression of the 1000 most variable genes identified in the murine data, or the imputed (t = 6) human data (Dermadi et al., 2020). The average of two trajectories in the murine datasets, and one trajectory in the human dermal dataset, identified as spanning from arterial to venous BEC was separated into equidistant bins and the mean gene expression was calculated for each bin and for each organ. The bins were subsequently annotated based on the mean expression as belonging to the subtypes described in the main text. UMAP calculations were done with umap (v.0.2.7.0) using imputed expression of the 1000 most variable genes. Heatmaps were generated using ComplexHeatmap (v.2.6.2), and violin plots were generated using ggplot2 (v.3.3.5) or GraphPad Prism (v.9.0.0). Both plot types display imputed gene expression (t = 2, murine data; t = 1, human data) without MNN-integration. Differential gene expression analysis comparing one subtype to all other cells in a sample was carried out using Seurat (v.4.0.5) *FindConservedMarkers* (mouse datasets) or *FindMarkers* (human dataset).

### Western blot analysis

Lungs from control and *Cldn5* iECKO mice were removed and snap frozen. Protein was obtained by mechanical dissociation in RIPA buffer supplemented with 50nM Na3VO4, Phosphatase inhibitor cocktail (Roche 04906837001) and Protease inhibitor cocktail (Roche, 04693116001). LDS sample buffer (Invitrogen, NP0007) and Sample Reducing Agent (Invitrogen, NP0009) were added to the samples and heated to 70°C for 10 minutes. Proteins were separated on Nu Page 4-12% Bis-Tris Gel (Invitrogen) in MOPS SDS Running buffer (Invitrogen, NP0001), transferred to PVDF membrane (Thermo scientific, 88518) in NuPAGE transfer buffer (Novex, NP006), 10% methanol and subsequently blocked with 5% BSA in Tris-buffered saline with Tween 20 (TBST) for 60 minutes. The immunoblots were analysed using primary antibodies incubated overnight at 4°C and secondary antibodies linked to horseradish peroxidase (HRP) (Cytiva) incubated for 1 hour at room temperature. After each step filters were washed four times with TBST. HRP signals were visualised by enhanced chemiluminescence (ECL) (Cytiva) (1:25000) and imaged with Chemidoc (Bio-Rad). Primary antibodies targeting GAPDH (Chemicon, MAB374), Cldn5 (ThermoFischer Scientific, 352500), ZO-1 (ThermoFischer Scientific, 402200), Occludin (ThermoFischer, 711500), VE-Cadherin (R&D Systems, AF1002), JAM-A (Martin-Padura et al., 1998), Cingilin (Cardellini et al., 1996) and ESAM (R&D Systems, AF2827) were used at a dilution of 1:1000.

### Quantitative PCR

Lungs from control and *Cldn5* iECKO mice were removed into RNAlater (ThermoFischer Scientific, AM7024). RNA was extracted and purified using RNeasy Plus kit (Qiagen). RNA concentrations were measured by Nanodrop spectrophotometer (ThermoFisher Scientific) and adjusted to equal concentration, followed by reverse transcription using iScript Adv cDNA Kit for RT-qPCR (Bio-Rad, 1725038). Real-time quantitative PCRs were performed on Bio-Rad real-time PCR machine using the Taqman assay with the following probes: *Gapdh* (Mm99999915_g1), *Cldn5* (Mm00727012_s1), *Cdh5*(Mm00486938_m1), *Tjp1* (Mm01320638_m1), *Ocln* (Mm00500912_m1), *F11r* (Mm00554113_m1), *Cgn* (Mm01263534_m1), *Esam* (Mm00518378_m1). The comparative Ct method was used to calculate fold differences.

### Electron microscopy

To study junction ultrastructure, control or *Cldn5* iECKO mice were anesthetized and perfused first with 10 mL HBSS and then 12 mL cold fixative (1% GA, 4% PFA in 0.1 M phosphate buffer) through the left ventricle. To study HRP penetrance, control or *Cldn5* iECKO mice were intravenously injected with HRP (Sigma Aldrich, 77332) (20mg / 25g body weight) and histamine (0.25mg / 25g body weight) before anaesthetization and perfusion after 10 minutes. Ears were then removed and placed in fixative for 30 minutes at 4°C before washing in PBS. Samples with HRP were treated with 0.05% DAB in PBS for 20 minutes and then with buffer containing H_2_O_2_ and 0.05% DAB for 30 minutes. Samples were post-fixed with 1% OsO_4_ + 1,5% C_6_FeK_4_N_6_ 2 h at +4°C before dehydration and LX112 resin infiltration. Post-staining was with C_12_H_10_O_14_Pb_3_. Samples without HRP were treated as above but devoid of DAB treatment and including 1% C_4_H_6_O_6_U (Uranyl acetate) staining. Samples with HRP were treated with 0.05% DAB in PBS for 20 minutes and then with buffer containing H_2_O_2_ and 0.05% DAB for 30 minutes. Samples were post-fixed with 1% OsO_4_ + 1,5% C_6_FeK_4_N_6_ for 2 hours at 4°C before dehydration and LX112 resin infiltration. Post-staining was with C_12_H_10_O_14_Pb_3_. Samples without HRP were treated as above but devoid of DAB treatment and including 1% C_4_H_6_O_6_U (Uranyl acetate) staining. Transmission electron microscopy (TEM) imaging and analysis was done at the Euro-BioImaging facility at Biocenter Oulu, Finland.

### Statistical analysis

Data are expressed as mean ± SD. The principal statistical test used was the Students’ t test. *p-values* given are from independent samples analyzed by two-tailed paired t tests. Rate of leakage was compared using linear regression and ANCOVA. Correlations were calculated using a Pearson’s correlation. All statistical analyses were conducted using GraphPad Prism. A *p-value* < 0.05 was considered statistically significant and significances indicated as p < 0.05 (*), p < 0.01 (**), and p < 0.001 (***). For animal experiments no statistical methods were used to predetermine sample size. The investigators were blinded to allocation during experiment and outcome assessment.

### Data availability

The murine ear skin data has been deposited in GEO under accession number (TBD). Further details regarding specifics of the analysis will be available upon reasonable request.

